# Decoding Phenotypic Variability in Osteogenesis Imperfecta: Zebrafish as a Model for Molecular and Ultrastructural Insights

**DOI:** 10.1101/2025.01.28.635270

**Authors:** Sophie Debaenst, Lauren Sahd, Caitlin Debaene, Tamara Jarayseh, Hanna De Saffel, Jan Willem Bek, Wouter Steyaert, Delphi Van Haver, Sara Dufour, Francis Impens, Paul Coucke, Adelbert De Clercq, Andy Willaert

## Abstract

Phenotypic variability is common in human diseases, even when the same genes are affected. In this study, three zebrafish models of Osteogenesis Imperfecta (OI) with dominant glycine substitutions in type I collagen genes (*col1a1a^mh13/+^*, *col1a1a^dc124/+^*, and *col1a2^mh15/+^*) were characterized for phenotypic severity and variability, using a newly developed standardized scoring system. Comprehensive analyses of the vertebral columns in these models revealed histological and ultrastructural differences that corresponded with phenotypic severity. Increasing skeletal severity correlated with a higher incidence of skeletal deformities and abnormalities. This, in turn, was associated with thinner bones and increased disorganization of collagen fibrils, fiber accumulation and mineralization, elastin deposits, and increased cell proliferation in the notochord and intervertebral ligament (IVL). Additionally, osteoblast function and bone regenerative capacity were increasingly compromised. These characteristics, combined with genetic information, have the potential to predict the severity of phenotypic outcomes in dominant forms of OI, caused by mutations in type I collagen. A remarkable intra-familial phenotypic variability in the *col1a2^mh15/+^*mutant holds potential for future approaches that could help in understanding the underlying mechanisms of this variability and the identification of modifier genes. Finally, through proteomics analysis three potential protein biomarkers (HSP47, Col8a1, and Bcan) were identified, that could serve as indicators of disease severity. These biomarkers not only have diagnostic value, but will allow stratification by OI type, have predictive value towards progression of the clinical presentation and will play a role in treatment guidance. Validation in human tissue samples will further reveal their clinical relevance.

**Significance Statement:** Phenotypic variability in human diseases, such as Osteogenesis Imperfecta (OI), remains poorly understood. Using zebrafish models with dominant glycine substitutions in type I collagen, this study links genetic mutations to phenotypic severity through standardized scoring and detailed ultrastructural and molecular analyses. Key findings include skeletal abnormalities, compromised osteoblast function, and intra-familial phenotypic variability, suggesting the role of modifier genes. Proteomics identified three potential biomarkers (HSP47, Col8a1, and Bcan) with diagnostic and prognostic value. These results provide critical insights into genotype-phenotype correlations, offering a foundation for personalized approaches to diagnosis, stratification, and treatment of OI and related disorders.

## Introduction

Osteogenesis Imperfecta (OI) is a rare genetic connective tissue disorder that occurs in approximately 1 in 10,000 births (1). Key characteristics of OI include bone fragility, frequent fractures, short stature, and various extra-skeletal manifestations such as blue sclerae, skin and joint laxity, and muscle weakness (2). This disorder primarily results from dominant mutations in the *COL1A1* (OMIM 120150) or *COL1A2* (OMIM 120160) genes (3), which encode type I collagen. However, recessive inheritance patterns have also been observed in patients carrying mutations in other genes involved in osteoblast differentiation, bone mineralization and collagen processing (4). The Online Mendelian Inheritance in Man (OMIM) database recognizes 22 types of OI based on the affected gene and their phenotypic characteristics (2). Several of these types are progressive and may worsen with age (5).

Phenotypic variability or differences in clinical presentation are well documented in OI patients, who exhibit varying phenotypes not only in the case of different genetic causes but even in the presence of identical mutations (6–9). In dominant types of OI, phenotypic variability can be determined by the nature of the specific mutations in the *COL1A1* and *COL1A2* genes, which are responsible for structural defects in type I collagen (2). Factors contributing to this variability include the specific collagen gene affected, the chemical properties of the substituted amino acid, the position of the mutation within the gene, or a combination of these factors (2,10). For instance, glycine substitutions within these genes lead to phenotypes ranging from mild to lethal, classified as OI types I to IV (11). In addition, a recent study analyzing *in vitro* cultures of bone cells has identified differentially expressed proteins, including proteoglycan, decorin, biglycan and osteonectin, that may influence the phenotypic variability in both lethal and non-lethal forms of OI (2). Proteomic analysis of fibroblast cells from three patients with lethal OI and three with non-lethal OI, all harboring glycine substitutions in the *COL1A1* or *COL1A2* genes, revealed a downregulation of the extracellular matrix protein decorin and an upregulation of fibrillin-1 in those with lethal OI (12). In another study by Marini *et al.* (13), bone proteome profiling in the Brtl murine model suggests that variations in extracellular matrix composition, even with identical collagen mutations, may contribute to the differing phenotypic outcomes. Nevertheless, further investigation is needed to unravel the factors determining phenotypic variability in dominant OI.

Zebrafish (*Danio rerio*) are well suited for studying skeletal diseases. Their small size, fully sequenced genome and large number of offspring, constitute core advantages (14). Furthermore, zebrafish can tolerate more severe skeletal deformities compared to rodent models, likely due to a reduced impact of gravity on their skeleton in their aquatic environment (15). These features have facilitated the generation and characterization of several mutant and transgenic zebrafish models that mimic OI (16–20). Unlike traditional rodent models, zebrafish exhibit an outbred nature that more closely mirrors the human genetic constitution, allowing for the study of phenotypic variability. Notably, zebrafish show a wide range of phenotypic variations of skeletal elements in response to environmental factors such as rearing density (21), highlighting that a variable genome can express vast variation under different environmental conditions. Consequently, mutant zebrafish that mimic human OI phenotypes, including the expression of phenotypic variation in the skeleton, are an interesting model to study the variable manifestations of this disease.

Zebrafish models of OI revealed significant variability of skeletal abnormalities as reviewed by Masierio *et al.* (22). For instance, Gistelinck *et al.* (23) found that zebrafish with different heterozygous glycine substitutions in type I collagen exhibited varying phenotypic severities, mainly affecting the vertebral column. Furthermore, *plod2*^-/-^ mutants showed vertebral compression, kyphosis, and scoliosis, with microCT imaging revealing ectopic bone formation and irregular bone structures within the vertebral column (19). Zebrafish knockouts for *crtap* and *p3h1*, which mimic human mutations, displayed overmodified collagen type I and a recessive OI phenotype characterized by vertebral deformities and fractures (3,20). Zebrafish models lacking *fkbp10a* showed reduced lysyl hydroxylation of type I collagen, resulting in skeletal variability, enlarged collagen fibrils and disturbed elastin layers (24).

Understanding the molecular mechanisms driving phenotypic variation in OI is crucial for developing therapeutic targets and precision medicine. Both environmental and genetic factors appear to play a significant role in skeletal phenotypic diversity, and it is yet to be discovered if one has a more prominent role than the other (21). Comprehensive phenotyping of zebrafish OI models is essential to link molecular changes to skeletal alterations. In addition, identifying molecular signatures can offer diagnostic value, enable stratification by OI type, predict disease progression, and guide treatment strategies.

This study investigates the range of skeletal phenotypes in sibling OI zebrafish with identical type I collagen mutations as well as the specific phenotypic differences in zebrafish OI models with different type I collagen mutations. To achieve this, a standardized scoring system for assessing skeletal disease in zebrafish models was established. This system enabled detailed characterization of phenotypic variability in three different OI zebrafish mutants: *col1a1a^mh13/+^*, *col1a1a^dc124/+^* and *col1a2^mh15/+^*. Additionally, comprehensive analyses were conducted to identify (ultra)structural differences and potential molecular biomarkers for phenotypic variability using histology combined with transmission electron microscopy and proteomic analysis, respectively.

We hypothesize that zebrafish models of OI with different mutations exhibit distinct skeletal phenotypic variability and ultrastructural differences. This variability may be reflected in specific molecular biomarkers, which could be identified through detailed molecular analysis.

## Results

### Phenotypic scoring reveals variable vertebral phenotypes across three different dominant OI zebrafish models

Three osteogenesis imperfecta (OI) zebrafish models with dominant glycine substitutions in the genes encoding type I collagen were selected to study variability in the vertebral column at both phenotypic and molecular levels; The *col1a1a^mh13/+^*, *col1a1a^dc124/+^* and *col1a2^mh15/+^*mutants exhibited mild, severe, and ‘moderate to very severe’ visual expressions of the OI phenotype, respectively (23). A novel scoring matrix, designed to enable reproducible skeletal phenotyping based on Alizarin Red S mineral-stained whole skeletons, was developed (see M&M and Table S1 and Dataset S1) and assessed in *col1a1a^mh13/+^(*n=13), *col1a1a^dc124/+^* (n= 15) and *col1a2^mh15/+^* (n= 11) mutants and their respective wild type siblings (n=8, n=13 and n=11 respectively). The deformity frequencies were determined as a percentage of the number of deformed vertebrae relative to the total number of vertebrae studied (Figure 1A). The *col1a1a^mh13/+^* mutants appeared similar to their wild-type counterparts with few deformities, except for notochord (NC) mineralization (9.4%), intervertebral ligament (IVL) mineralization (25.8%), detachments (15.7%) and scoliosis (21.1%). The *col1a1a^dc124/+^* mutants exhibited a more defined phenotype characterized by NC mineralization (68.1%), IVL mineralization (80.2%), fractures (28.5%), detachments (93.6%), fused vertebrae (12.8%) and compressed vertebrae (15.8%) and scoliosis (22.1%). Similarly, *col1a2^mh15/+^* mutants were distinctly characterized by extensive additional mineralization of the NC (71.5%) and IVL (69.7%), coupled with prominent scoliosis (47.7%), detachments (88.8%), as well as fused vertebrae (18.8%), compressed vertebrae (12.3%) and fractures (12.6%).

**Figure 1.**
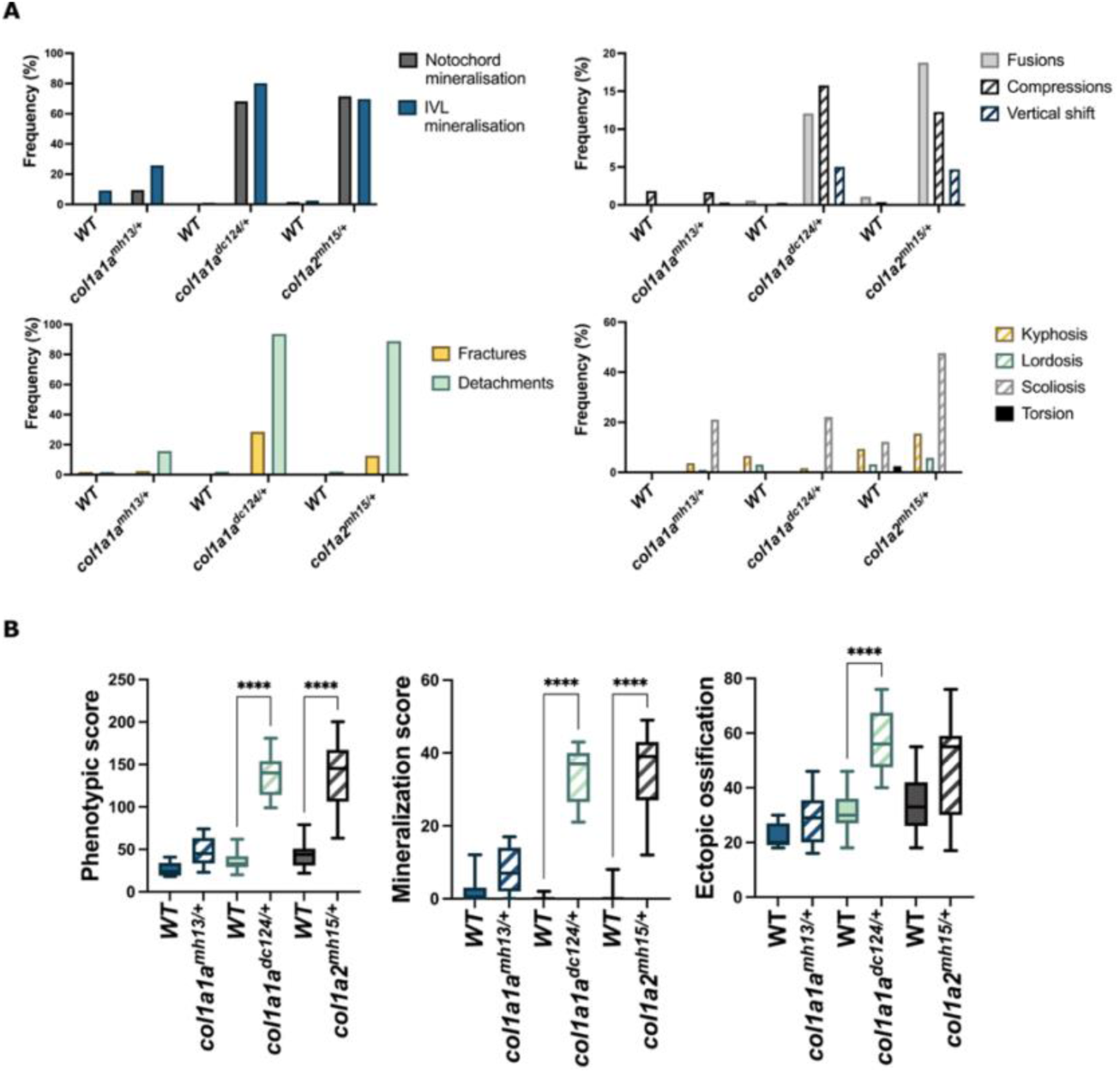
The frequency of deformities in each mutant model (A) and comparison of the phenotypic scores between mutants and respective sibling wild-type controls (B). Frequencies were calculated as the total number of vertebrae affected relative to the total number of vertebrae analysed. **** indicates p< 0.001.

A phenotypic score, derived from the total number of deformities indicated in the scoring matrix (see M&M), was calculated in each zebrafish individual. All three mutants showed a higher phenotypic score compared to their respective wild-type controls (Figure 1B). However, statistical significance was observed only in the *col1a1a^dc124/+^* and *col1a2^mh15/+^*mutants (p<0.0001; mean: 135.54 ± 25.21 and 135.27 ± 43 respectively), indicating a comparatively milder phenotype in the *col1a1a^mh13/+^* mutants (p= 0.21; mean 48.92 ± 17.17) (Figure 1B). The *col1a1a^mh13/+^* mutants did not exhibit significant differences in either the mineralization score (p= 0.33; mean: 8.23 ± 6.27), assessed by the total number of vertebrae displaying IVL and NC mineralization, or the ectopic ossification score, which is based on the extra mineralized bone on the vertebral body (p= 0.56; mean: 29.08 ± 8.97), compared to their wild-type control (Figure 1B). In contrast, *col1a1a^dc124/+^*mutants, showed a significant increase in both the mineralization score (p<0.0001; mean: 34 ± 7.86) and ectopic ossification score (p<0.0001; mean: 57.08 ± 11.46) compared to their wild-type controls. The *col1a2^mh15/+^*mutants showed a significant increase only in the mineralization score (p<0.001; mean: 35.55 ± 11.1) compared to their wild-type controls. Interestingly, the *col1a2^mh15/+^* mutants exhibited a wide data range in all three total scores (mean phenotypic score: 135.27 ± 43; mean mineralization score: 35.55 ± 11.1; mean ectopic ossification score: 46.64 ± 19.76) suggesting substantial phenotypic intra-variability within this mutant group. Additionally, although not further investigated in detail, a collapse of the nasal structures and their strong deformation or even absence was observed in the *col1a1a^dc124/+^* and severe forms of *col1a2^mh15/+^* fish, but not in the mild *col1a1a^mh13/+^*fish. Representative images of each mutant and their respective wild-type controls are depicted in Figure 2, showing the mild phenotype of *col1a1a^mh13/+^,* the defined severe phenotype of *col1a1a^dc124/+^* and the variable phenotype of *col1a2^mh15/+^* which ranges from moderate to very severe.

**Figure 2.**
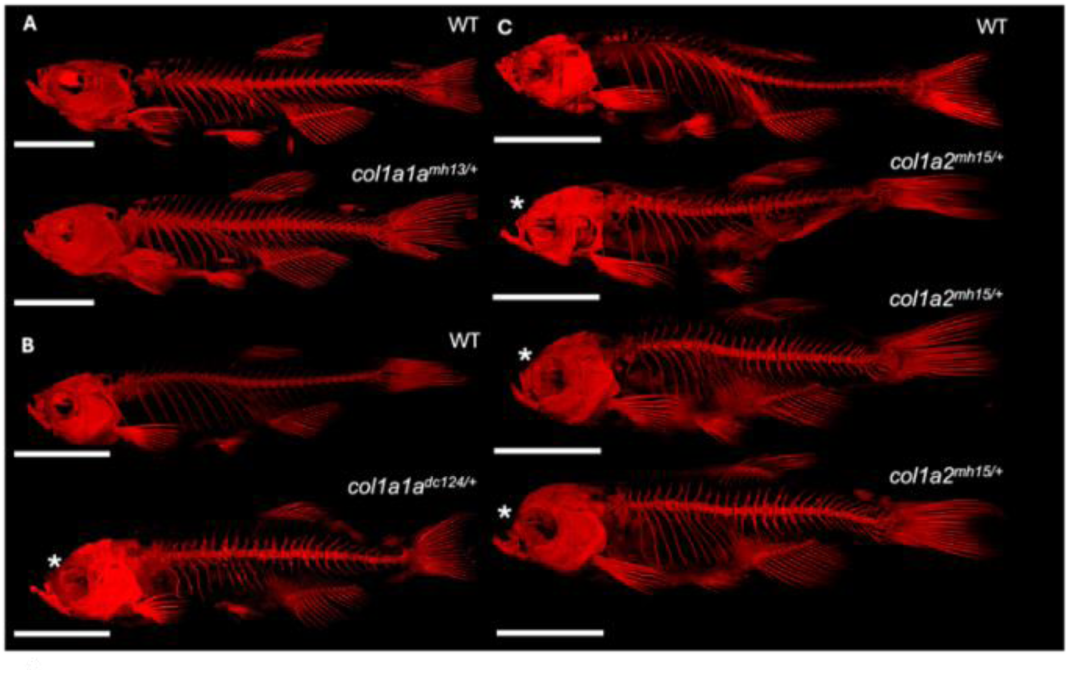
Alizarin Red S mineral-stained whole skeletons of the *col1a1a^mh13/+^* (A), *col1a1a^dc124/+^* (B) and *col1a2^mh15/+^* (C) model and respective wild-type siblings. Mutants for *col1a2^mh15/+^* (C) showing the range of phenotypic variability, with up to down: Wild-type - moderate representative - severe representative – very severe representative. Skeletons stained with Alizarin Red S for mineralized tissues and imaged using its autofluorescent properties. Asterix indicates deformed nasal structures. Scale bar = 5mm.

### Fin regeneration is variably affected in dominant OI zebrafish

To assess and compare bone formation and regeneration capacities in the three collagen mutants, a fin regeneration experiment was conducted. Immature osteoblasts (fin ray-forming cells, also referred to as scleroblasts) were visualized at 8 days post-amputation (dpa) using the transgenic *osx:*Kaede zebrafish line. Photoconversion from green to red fluorescence was performed at 0 dpa, allowing for the quantification of the newly differentiated green-colored *osx*-positive cells at 8 dpa. Additionally, the morphology of the new fully regenerated fins was evaluated at 21 dpa. The 8 dpa and 21 dpa time points in zebrafish fin regeneration were selected to capture key stages of regeneration. At 8 dpa, active growth and proliferation were ongoing, whereas at 21 dpa, final morphology and patterning could be evaluated.

The analysis revealed no significant differences in the length of the newly formed fin rays between *col1a1a^mh13/+^* zebrafish (n=5) (mean: 1.31 ± 0.12) and their respective controls (n=4) (mean: 1.27 ± 0.25) at 8 dpa (p= 0.76) (Figure 3A). Morphologically, both wild-type controls (mean: 1.73 ± 0.27) and *col1a1a^mh13/+^* mutants (mean: 1.59 ± 0.15) displayed a similar, typical fan-shaped tail fin with no statistical difference in tail width (p= 0.32) (Figure 3B). In contrast, *col1a1a^dc124/+^* fish (n=5) (mean: 1.16 ± 0.14) exhibited a significant reduction (p<0.001) in the length of newly formed fins at 8 dpa compared to their respective controls (n=5) (mean: 1.83 ± 0.05). Additionally, by 21 dpa, the regenerated fins were significantly narrower (mean: 1.18 ± 0.27) than the controls (mean 1.60 ± 0.15) and exhibited disturbances in the typical fan shape (p<0.05) (Figure 3A-C). While some *col1a2^mh15/+^* individuals exhibited reduced fin ray length at 8 dpa, no significant differences were observed in fin ray length between *col1a2^mh15/+^* zebrafish (n=4) (mean: 1.28 ± 0.43) and their respective controls (n=5) (mean: 1.29 ± 0.33) (p= 0.97). Additionally, at 21 dpa, some individuals within the *col1a2^mh15/+^*group (mean: 1.31 ± 0.53) displayed abnormalities in fin morphology compared to their respective controls (mean: 1.66 ± 0.18), although not reaching statistical significance (p= 0.26) (Figure 3A-D). This correlated with the high phenotypic variability observed in the vertebral column of this mutant line.

**Figure 3.**
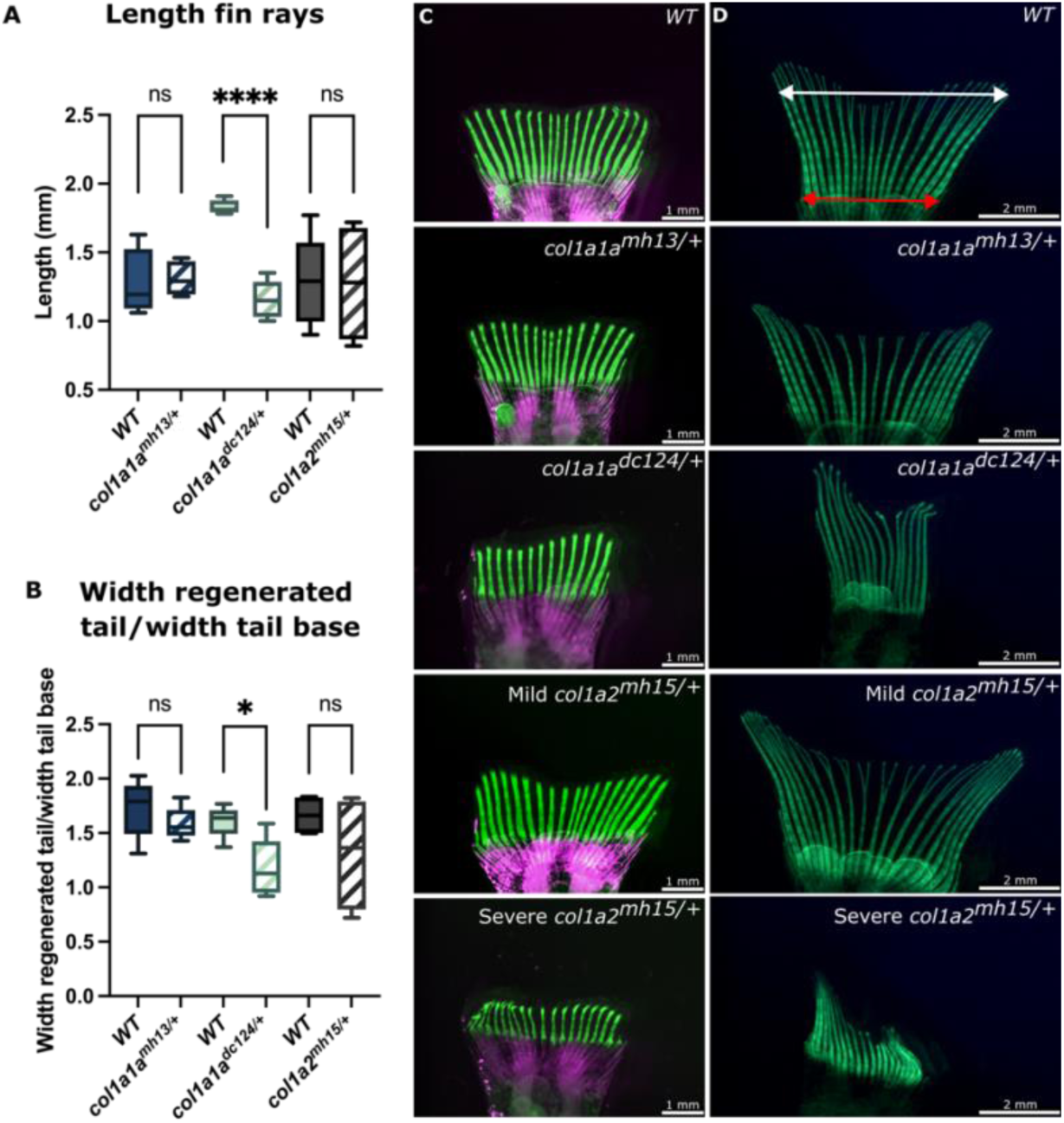
Fin regeneration in *col1a1a^mh13/+^*, *col1a1a^dc124/+^*, and *col1a2^mh15/+^* mutants and their respective wild-type siblings. The *osx:*Kaede transgenic line was used to image early osteoblasts following photoconversion at 0 dpa. (A) At 8 dpa, the length of newly generated fins rays (in total 12 rays per fish, specifically ray 2-7 and 11-16, from dorsal to ventral) was quantified in the green channel, starting at the cut site. (B) Morphological analysis of fully regenerated fins, based on tail width normalized to the tail base. (C) Images of the regenerating fins at 8dpa in WT, *col1a1a^mh13/+^*, *col1a1a^dc124/+^*, mild *col1a2^mh15/+^* and severe *col1a2^mh15/+^*. (D) Images of the fully regenerated fins at 21 dpa in in WT, *col1a1a^mh13/+^*, *col1a1a^dc124/+^*, mild *col1a2^mh15/+^* and severe *col1a2^mh15/+^*. The white double arrow marks the maximum tail width measured, while the red double arrow represents the tail base.

### Histological and ultrastructural analysis reveals a correlation between collagen organization and notochord mineralization with phenotypic severity in dominant OI zebrafish models

A comprehensive histological analysis of the vertebrae in the three OI models was conducted to gain deeper insights into the pathological traits underlying the various vertebral deformities observed in whole-mount stained vertebral columns for mineralized tissues. Morphologically, the *col1a1a^mh13/+^*mutants displayed a mild phenotype characterized by a few deformities, including scoliosis, occasional compressed vertebrae, and minor ectopic mineralization. The presence of these mild skeletal deformities was reflected in their mild external phenotypes. Therefore, *col1a1a^mh13/+^* zebrafish and wild type siblings (n = 3 for each genotype) were selected at random for histological analyses. Serial sections of the vertebral column showed vertebral bodies with normal vertebral centra, presenting regular autocentra, as well as normal notochord tissue, recognizable as the septum present in the intervertebral region (Figure 4A-C).

**Figure 4.**
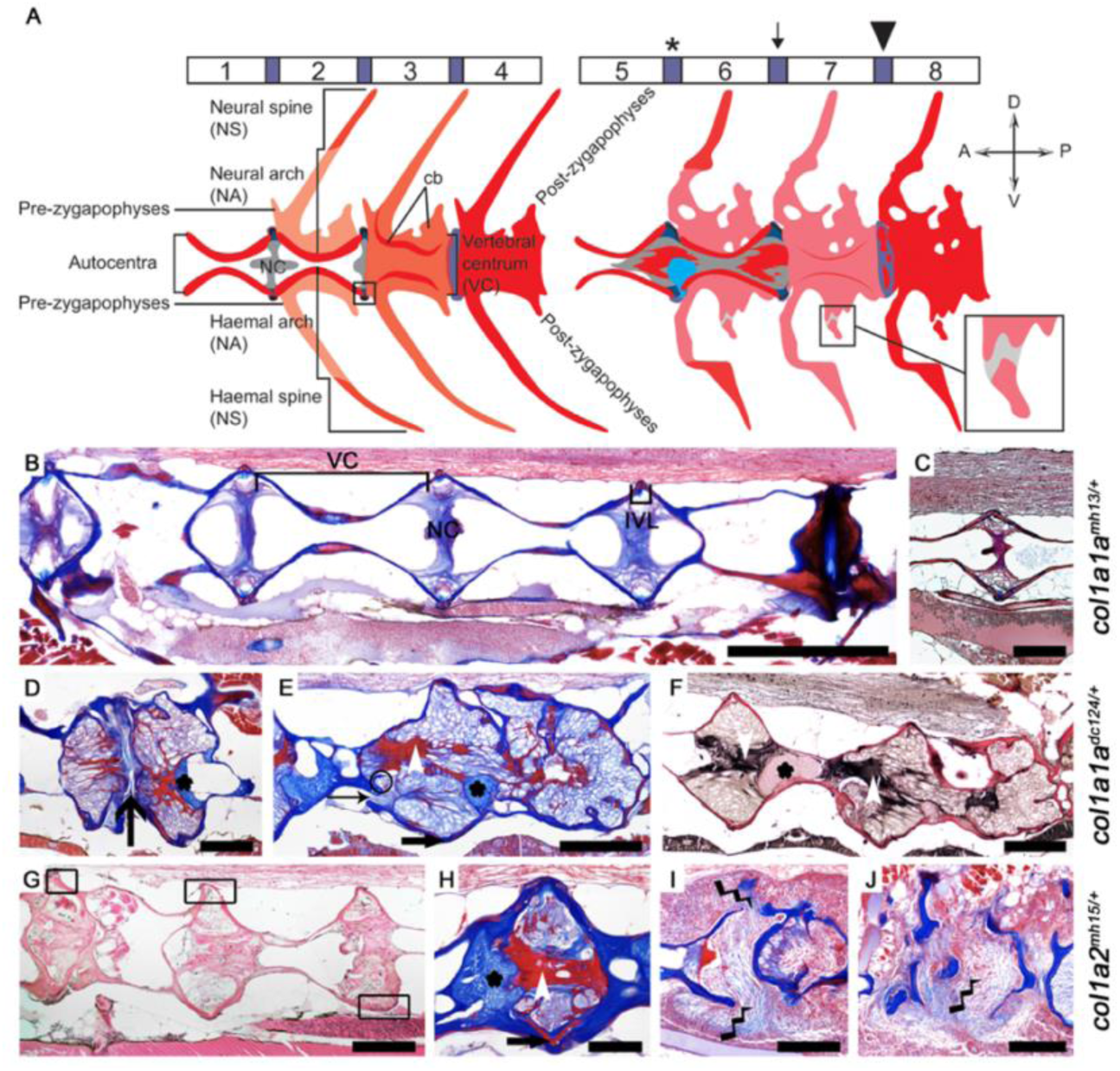
Histology of the vertebral column of OI mutants. Schematic overview of wild type (1–4) and mutant vertebrae (5–8) (A). All structures that were separately scored on whole-mount stained animals are indicated. The purple blocks at the top indicate the intervertebral disc regions. Vertebrae 1 and 5 show only the autocentra as seen in sections. Vertebrae 2 and 6 show autocentra with the dorsal and skeletal elements, vertebrae 3 and 7 show vertebrae as observed with fall-through white light and vertebrae 4 and 8 show vertebrae as observed with autofluorescence. The black box on the ventral side between vertebrae 2 and 3 shows an intervertebral ligament which is shown enlarged in Figure 6A. The askterisk shows an intervertebral disk with fibrous tissue at its center (red) and cartilage inside the notochord (blue), notice that the cartilage can herniate from the notochord. The black arrow shows only the presence of fibrous tissue (red) in the notochord, and the black box on the ventral side of vertebra 7 shows a seemingly detached piece of trabecular bone, but notice the non-stained collagen fibers connecting this detached bone (enlarged on the right). The arrowhead shows patches of mineral in the intervertebral ligament as observed with autofluorescence. Abbreviations: A, anterior, cb, cancellous bone, D, dorsal, NC, notochord, P, posterior, V, ventral. Images of serial sections of *col1a1a^mh13/+^* (B-C), *col1a1a^dc124/+^*(D-F) and *col1a2^mh15/+^* (G-J) mutants. Symbols: line arrow, location of compression fracture, small black arrow, detachment of bone, white arrowhead, fibrous material inside the notochord, black asterisks, cartilage in the notochord, bock arrow, IVL mineralization, lightning bolts, fibrotic tissue replacing the notochord and bone of the vertebral column. Abbreviations: IVL, intervertebral ligament, NC, septum of the notochord, VC, vertebral centrum. Staining: AZAN, B, D-E, H-J; Verhoeff-Van Gieson, C, F; Hematoxylin & Eosin (H&E), G. Scale bars: A = 1 mm, C-G, I-J =500 µm, H = 250 µm.

The *col1a1a^dc124/+^* mutant displayed frequent fractures, compressed vertebrae and generally deformed vertebral bodies, presenting a severe external phenotype. As most *col1a1a^dc124/+^* mutant zebrafish are severely affected, random mutant and wild type siblings were selected for histological analyses (n = 3 for each genotype). Serial sections revealed vertebrae where the autocentra and cancellous bone appeared completely collapsed (Figure 4D; line arrow) resulting in juxtaposed intervertebral disks. In some cases, the collapse of the vertebral centrum was less severe, occurring only on the dorsal side (Figure 4E-F). The notochord septum appeared highly vacuolized and contained fibrous tissue at its central part in the middle of the vertebral disc (Figure 4A, E-F; white arrowhead). This notochord phenotype was present in both less severely and severely collapsed vertebrae, often in combination with the presence of fiber cartilage (Figure 4A, D-F; asterisks). A detached autocentrum was observed to be surrounded by fibrous tissue (Figure 4E; line arrow). The presence of osteoclasts (Figure 4E; circle) suggested bone remodeling surrounding the detached autocentrum. Additionally, Von Gieson’s staining highlighted fiber accumulation in the intervertebral space, which likely correlates with the prominent IVL and NC mineralization, previously observed with AR staining (Figure 4 C-E).

The skeletal morphology of *col1a2^mh15/+^* mutants exhibited a wide phenotypic range which was reflected in their external phenotype. Alongside wild type siblings, a moderately, severely and very severely affected mutant zebrafish were selected for histological analyses. Serial sections of the moderately affected mutants showed vacuolized notochord septa with low amounts of fibrous tissue and fiber cartilage, but recognizable intervertebral discs. The vertebrae with autocentra were also identifiable (compare Figure 4A scheme with Figure 4K); however, the vertebral end plates were deformed (Figure 4G; black boxes) distorting the intervertebral ligament. Similar to *col1a1a^dc124/+^*zebrafish, severely affected *col1a2^mh15/+^*mutants showed accumulated fibrous tissue in the center of the notochord septum (Figure 4H; white arrowhead), which could co-occur with fiber cartilage (Figure 4H; asterisks). In the most severely affected animals, complete detachment of the autocentra of the vertebra could be observed (Figure 4I-J). Parasagittal serial sections through the entire vertebral column showed no connection of bony or notochord tissue. Instead, fiber-rich tissue consisting of fibroblast-like cells occupied the space where the bone and notochord of the vertebral centrum were expected (Figure 4I-J; lightning bolt).

Observing the mineral in the intervertebral ligament, a skeletal morphological characteristic of both the *col1a1a^dc124/+^* and *col1a2^mh15/+^* mutants, was impossible on demineralized histological sections. In AZAN-stained sections collagen fibers stain blue but can stain red when they were mineralized before. Therefore, the presence of red-stained collagen fibers at the level of the intervertebral ligament likely indicates locations of mineralization of the intervertebral ligament in both mutants (Figure 4E, F; block arrow)

To investigate whether the mineralized core observed in the notochord of whole-mount stained *col1a1a^dc124/+^* and *col1a2^mh15/+^* mutants, corresponds to the fibrous tissue in the notochord observed in serial histological sections, non-decalcified sections were prepared from GMA-embedded vertebrae. Von Kossa staining, which specifically stains mineralized tissue, was used and validated with alizarin red S staining of directly subsequent sections. No staining of mineralized tissues was observed in the vertebral discs of wild type zebrafish (Figure 5A-B). Both methods stained the fibrous tissue in the vertebral disc of *col1a1a^dc124/+^* (Figure 5C-D; black arrowhead) and *col1a2^mh15^* (Figure 5E-F, black arrowhead) mutants, confirming that the fibrous tissue in serial histological sections was indeed the same as the mineralized core of the notochord observed in whole mount-stained specimens of these mutants.

**Figure 5.**
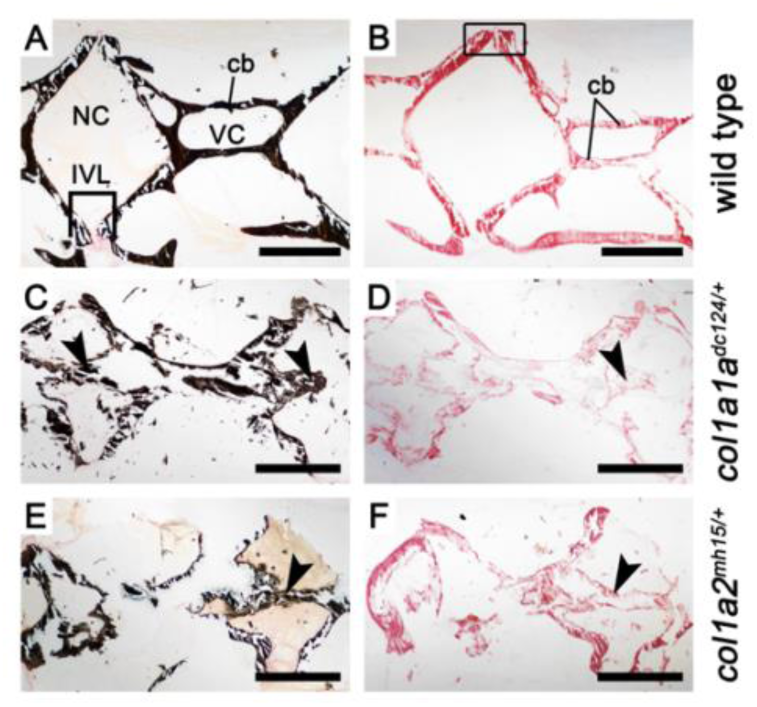
Mineralization of the notochord in non-decalcified GMA-sections. (A) Von Kossa-stained section of a wild type zebrafish, where mineralized tissue (phosphate) is stained black and unmineralized tissue is stained pink. There is no mineralized tissue in the intervertebral space. B: Alizarin red stained section of a wild type zebrafish, mineralized tissue (calcium) is stained red. The black box shows the vertebral endplates of two adjacent vertebrae. (C-D) A representative of the *col1a1a^dc124/+^* mutant with serial sections stained with Von Kossa (C) and alizarin red (D) showing mineralization of the notochord as indicated by the black arrowhead. (E-F) A representative of the *col1a2^mh15/+^* mutant with serial sections stained with Von Kossa (E) and alizarin red (F) showing mineralization of the notochord as indicated by the black arrowhead. Abbreviations: cb, cancellous bone, IVL; Intervertebral ligament, VC, vertebral centrum. Scale bars = 100 µm.

The ultrastructural characteristics of the intervertebral ligament were further investigated in the OI zebrafish models. Both for *col1a1a^mh13/+^* and *col1a1a^dc124/+^*, six animals and six WT sibling controls were chosen at random. For *col1a2^mh15/+^* again a range of six moderately, severely and very severely affected animals were selected. Transmission electron microscopic (TEM) was performed at the level of the intervertebral ligament and vertebral end plates. The vertebral ligament consists of well-described layers, from the inside of the notochord to the outside: a collagen type II layer and an elastin layer (together forming the notochord sheath), a mature collagen layer of compact collagen type I fibers and an immature collagen layer of loosely packed collagen type I layers (Figure 6; schematic). The vertebral endplates consist of intramembranous bone composed of the autocentra which are the continuation of the mature collagen layer of the intervertebral ligament into the bone as Sharpey’s fibers and the intramembranous bone that forms around the autocentra (Figure 6 schematic, with the dashed line indicating the separation between the autocentrum and bone).

**Figure 6.**
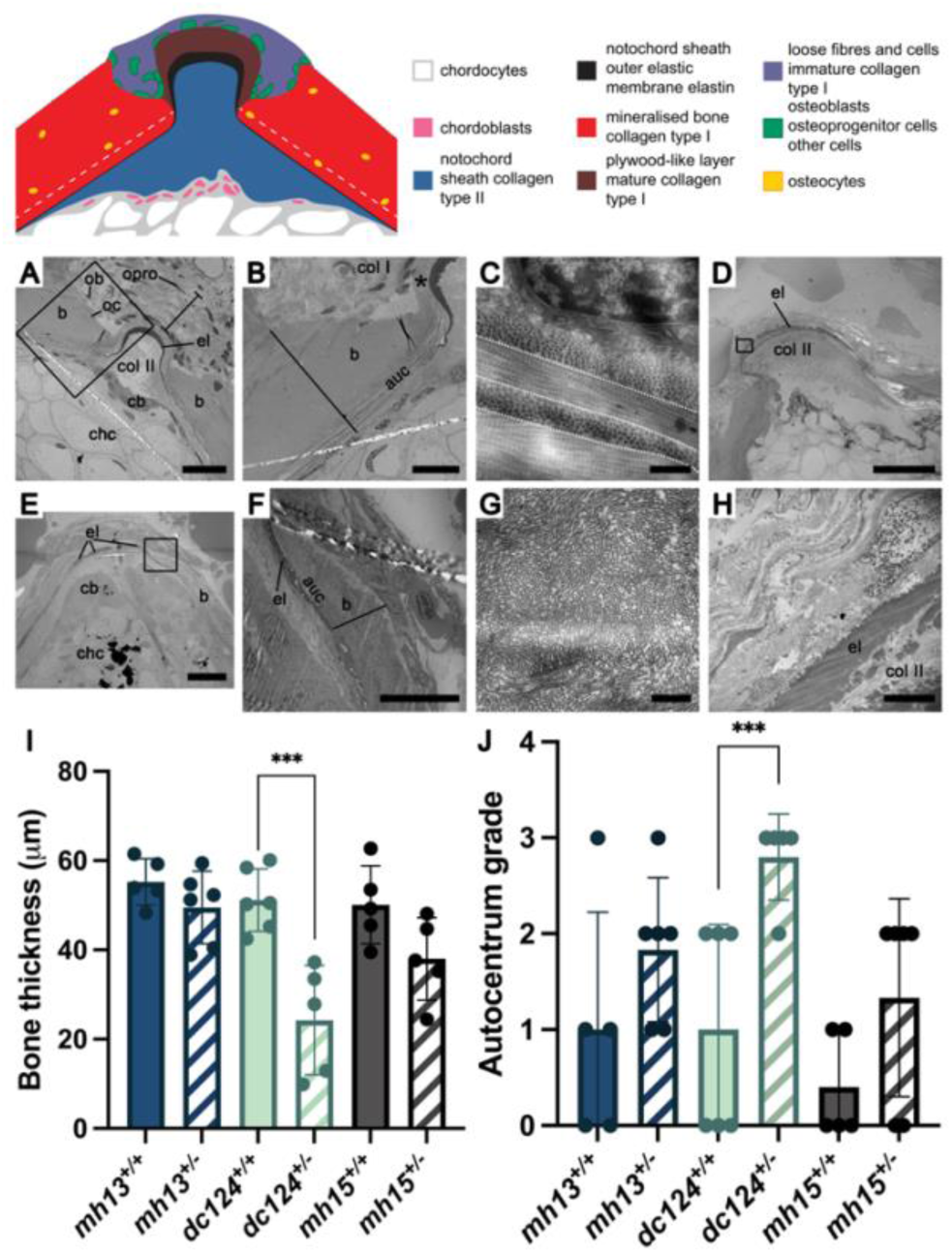
Transmission electron microscopy (TEM) analysis of OI zebrafish models. Schematic represents the intervertebral ligament and the vertebral endplates of two consecutive vertebrae. The schematic legenda explains the cells and layers indicated in different colors. The white dotted line indicates the transition between bony autocentrum and the bone of the vertebral centrum. (A-D) show images of WT siblings, (E-G) show images of *col1a1a^dc124/+^*, H shows an image of *col1a2^mh15/+^*. Black boxes in A and E show enlarged images in B and F, respectively. Black lines in (B) and (F) show bone thickness and location where bone thickness was measured. Asterisk in (B) shows the mature collagen layer. The image in (B) shows a grade 0 autocentrum and in F shows a grade 3 autocentrum. (C, G) show high magnification images of the mature collagen layer. Black box in (D) (grade 2 elastin) shows enlarged images in (H), notice the thinner collagen type II fibers interspersed with elastin. Abbreviations: auc, autocentrum, b, bone, cb, chordoblasts, chc, chordocytes, col I, collagen type I, col II, collagen type II, el, elastin, ob, osteoblast, oc, osteocyte, opro, osteoprogenitor cell. Scale bars: (A, E) = 20 µm, (B, F) = 10 µm, (C, G) = 500 nm, (D) = 40 µm, (H) = 2 µm.

During the TEM analysis a number of detailed features of the IVL were assessed including the number of different cell types, thickness of collagen layers and grade of different key structures (Table 2, Table S2, Figure 6A schematic). The number of chordoblasts, cells on the mature collagen, cells on the bone of both vertebral end plates lining the IVL and the total number of cells in the IVL but outside the notochord were counted during live TEM imaging. Counts were repeated in six mutant and wild type siblings per genotype, averages calculated for mutant and wild type siblings, and compared using a One-Way ANOVA per genotype. Chordoblast numbers were on average higher in both *col1a1a^dc124/+^*and *col1a2^mh15/+^* mutants and similar in *col1a1a^mh13/+^* mutants compared to their WT siblings. The total number of cells of the intervertebral ligament were on average higher in *col1a1a^mh13/+^* and *col1a2^mh15/+^*mutants but similar in *col1a1a^dc124/+^*mutants compared to their WT siblings. Osteoblasts on mature collagen and bone were on average higher in *col1a1a^mh13/+^* mutants, respectively higher and lower in *col1a1a^dc124/+^* and respectively higher and similar in *col1a2^mh15/+^* mutants compared to their WT siblings.

Collagen type I-layer thickness and bone thickness were measured on TEM images. Measures in FIJI were repeated in six mutant and wild type siblings per genotype, averages calculated for mutant and wild type siblings, and compared using a One-Way ANOVA per genotype. The mature collagen layer consists of multiple layers of collagen type I fibers that alternate in direction from longitudinal (parallel to the middle line of the animal) to circumferential (perpendicular to the middle line of the animal) creating a plywood-like structure. The thickness of the mature collagen layer was, on average, similar in *col1a1a^mh13/+^* mutants compared to their WT siblings. The number of alternating layers was higher in these mutants compared to WT siblings. The similar average thickness of the mature collagen layer in both mutants and WT could be attributed to the thinner circumferentially oriented collagen type I layers in the mutants. In the *col1a1a^dc124/+^*mutants, the mature collagen layer was, on average, much thicker compared to their WT siblings. This increase in thickness could be explained by both a higher number of disorganized alternating layers (Figure 6G) and thicker longitudinally and circumferentially oriented collagen type I layers. Similarly, in *col1a2^mh15/+^*mutants, the mature collagen layer was, on average, thicker compared to their WT siblings. This could be due to a higher number of alternating layers even though both the longitudinal and the circumferential layers were, on average, thinner in the mutant animals. The bone was thinner in all OI zebrafish mutants (Figure 6I) compared to their WT siblings, but it was only significantly thinner in the *col1a1a^dc124/+^*mutants (Figure 6A-B for WT and 6E-F for mutant *col1a1a^dc124/+^*)

The intervertebral ligament (IVL) angle, a measure of symmetry in the IVL was assessed using TEM images. On average, the angle was higher in both *col1a1a^mh13/+^* and *col1a2^mh15/+^* mutants and similar in *col1a1a^dc124/+^* mutants compered to their WT siblings. The higher angle value indicated at least a mildly increased deformation of the IVL in both *col1a1a^mh13/+^* and *col1a2^mh15/+^*mutants. Grades were scored during live TEM imaging and repeated in six mutant and wild type siblings per genotype. The average grade was calculated and compared using a One-Way ANOVA per genotype. For correct biological interpretation when comparing the average grade, the value was rounded to the closest whole value. Both the elastin grade (Figure 6D, H) and ER grade were, on average, higher in all OI zebrafish mutants compared to their WT siblings. The autocentrum grade was also higher in all OI zebrafish mutants compared to their WT siblings, with a significantly higher grade observed in *col1a1a^dc124/+^* mutants (Figure 6J). Figure 6B shows a WT sibling with a recognizable autocentrum bony layer where the plywood-like pattern of the mature collagen layer in the IVL continues into the autocentrum of the bone. In contrast, this plywood-like organization is not visible in Figure 6F (*col1a1a^dc124/+^*mutant) where the autocentrum is difficult to distinguish.

### Proteomic Analysis of the Vertebral Column Uncovers Differential Expression of Matrisome-Associated Proteins in Severely Affected OI Zebrafish Models

To identify molecular signatures associated with the distinct outcomes in the three different OI zebrafish models, proteins were extracted from the vertebral column (excluding the Weberian apparatus and the last four vertebrae) of each mutant (*col1a1a^mh13/+^* (n=4) *col1a1a^dc124/+^* (n=5) and *col1a2^mh15/+^* (n=5)) and their matching WT siblings (n= 4,5 and 4 respectively). For the *col1a2^mh15/+^* mutant, zebrafish exhibiting phenotypes on the more severe end of the spectrum were selected to ensure uniformity. The extracted proteins were digested with trypsin, and the resulting peptide mixtures were analyzed using liquid chromatography-tandem mass spectrometry (LC-MS/MS). In total, we identified 3,346 proteins (Dataset S3) of which 2,070 proteins were quantified (Dataset S4). Protein expression levels were compared between each mutant and its respective WT sibling controls and significant differences were visualized in a Volcano plot (Figure 7, Dataset S5). We investigated which of these proteins are part of the zebrafish matrisome, which encompasses the entire repertoire of known extracellular matrix (ECM) proteins (25). These proteins are categorized into three groups: i) core matrisome (proteoglycan, collagen and ECM glycoproteins) ii) matrisome-associated proteins (secreted factors, ECM-affiliated proteins and ECM regulators), and iii) putative matrisome associated proteins.

**Figure 7.**
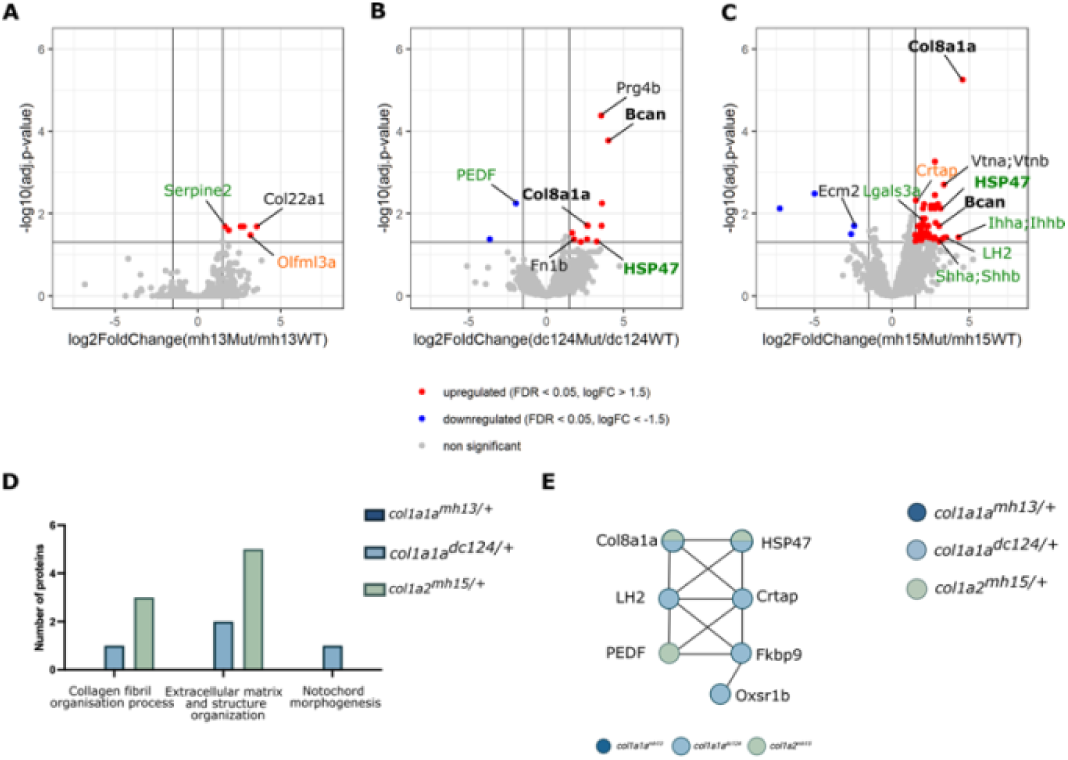
Volcano plots visualizing differentially expressed proteins in the (A) *col1a1a^mh13/+^*, (B) *col1a1a^dc124/+^* and (C) *col1a2^mh15/+^* mutants compared to their respective controls. Proteins significantly upregulated in the mutants are depicted in red, while downregulated proteins are shown in blue. For each protein, log2 (mutant/control) fold change values are shown on the X-axis, while the -log10 adjusted p-values indicating statistical significance (with false discovery rate (FDR) < 0.05) are shown on the Y-axis. In addition, protein labels of core matrisome proteins are depicted in black, matrisome-associated proteins in green and proteins with a putative matrisome association in orange. Proteins shared between the two severe mutants *col1a1a^dc124/+^*and *col1a2^mh15/+^* are indicated in bold. (D) Gene ontology analysis for biological processes, visualizing the number of differentially expressed proteins involved in three bone-related biological processes for each mutant. (E) Protein-interaction analysis, showing protein-interactions between the differentially expressed proteins in the mutants with severe phenotypes.

In the *col1a1a^mh13/+^* mutants, six proteins were significantly upregulated (Table S3, Figure 7A). Among these, three proteins, Col22a1 (core matrisome collagen), Serpine2 (matrisome-associated ECM regulator) and Olfml3a (putative matrisome association) belong to the zebrafish matrisome. The *col1a1a^dc124/+^* mutants showed significant upregulation of 11 proteins and downregulation of two proteins. Six of these proteins are part of the zebrafish matrisome, with four belonging to the core matrisome (Fn1b, Col8a1a, Bcan and Prg4b), and two being matrisome-associated ECM regulators (HSP47 and PEDF) (Table S3, Figure 7B). In *col1a2^mh15/+^* mutants, 61 proteins were significantly upregulated, while four proteins were downregulated. Comparative analysis of the differentially expressed proteins with the matrisome revealed four core matrisome proteins (Bcan, Col8a1a, Ecm2 and Vtna;Vtnb), five matrisome-associated proteins (Ihha;Ihhb, Lgals3b, LH2, HSP47 and Shha;Shhb) and one protein with putative matrisome association (Crtap) (Table S3, Figure 7C). Interestingly, three of the core matrix or matrix-associated proteins were found to be upregulated in both of the more severely affected mutants, *col1a1a^dc124/+^* and *col1a2^mh15/+^* (Bcan, Col8a1a and HSP47), while no commonly differentially expressed proteins were found between these two mutants and the milder *col1a1a^mh13/+^* mutant.

Gene Ontology analysis for bone-related biological processes in the three different mutants revealed that one of the differentially abundant proteins in the *col1a1a^dc124/+^* mutant (HSP47) and three in the *col1a2^mh15/+^* mutant (Crtap, LH2 and HSP47) are involved in the collagen fibril organization process. Two differentially abundant proteins in *col1a1a^dc124/+^* (Col8a1a and HSP47) and five proteins in *col1a2^mh15/+^* (Col8a1a, Crtap, Ecm2, LH2 and HSP47) were involved in extracellular matrix and structure organization (Figure 7D), and one differentially expressed protein in *col1a1a^dc124/^*+ mutants was linked to notochord morphogenesis (Col8a1a). A protein interaction analysis using STRINGdb (v11.0) identified interactions between two of the three matrisome-associated proteins that were upregulated in both the *col1a1a^dc124/+^* and *col1a2^mh15/+^* mutants but not in the *col1a1a^mh13/+^* mutants (Col8a1a and HSP47) (Figure 7E). Furthermore, other differentially expressed proteins, including LH2, Crtap, Fkbp9 and Oxsr1b in *col1a1a^dc124/+^* mutants and *Serpinf1* in *col1a2^mh15/+^* mutants, also interact at the protein level.

## Discussion

Osteogenesis imperfecta (OI) is a rare genetic connective tissue disorder characterized by substantial phenotypic variability, complicating its classification and genotype-phenotype correlation (4,13,26). Clinical variability occurs even among patients with identical mutations, making it difficult to predict outcomes based solely on genetic data (6–8, 11,27–29). Zebrafish have emerged as a valuable model organism for studying OI and its variable presentation, particularly for dominant forms of OI linked to type I collagenopathies (23). This study investigated three zebrafish OI mutants, *col1a1a^mh13/+^*, *col1a1a^dc124/+^* and *col1a2^mh15/+^*, all carrying heterozygous glycine substitutions in type I collagen and presenting with different phenotypic severities.

Zebrafish *col1a1a^mh13/+^* mutants exhibited a mild phenotype, closely resembling wild-type zebrafish. They presented minor skeletal deformities including subtle notochord mineralization defects, occasional vertebral compressions, and scoliosis, consistent with findings by Gistelinck *et al*. (23). Fin regeneration experiments revealed no differences in fin length at 8 days post-fertilization (dpa) or morphology at 21 dpa between *col1a1a^mh13/+^*mutants and wild-type siblings, indicating nearly intact osteoblast activity and bone formation. The skeletal morphological results were confirmed by the histological analyses which showed well-formed bony structures and nearly normal notochord characteristics. TEM analyses highlighted subtle changes in collagen layer organization both in number of fibers and their arrangement within the layers. The higher number of osteoblasts, exhibiting increased endoplasmic reticulum (ER)-stress, in the intervertebral ligament could indicate increased cellular retention of mutant collagen type I. The morphological, structural and ultrastructural analyses underscored the mild impact of the mutation. These results further confirm that the *col1a1a^mh13/+^* mutant carry qualitative alterations of collagen type I (22). Overall, this mutant tends to correspond to patients with the mildest OI type within the category of qualitative collagen type 1 defects (OI type IV) (22,23). It is challenging to contextualize results from this mild OI zebrafish model due to the limited scientific literature on mild phenotypes. Severe OI zebrafish models are more commonly studied because their phenotypic recognition is easier, and experimental results are clearer. Mild models may present confounding factors such as housing conditions and breeding programs. However, this study demonstrates that detailed investigations on mild OI zebrafish models are possible, revealing specific mild changes of collagen fiber organization and cell dynamics.

The Chihuahua mutant (*col1a1a^dc124/+^*), a well-established zebrafish model for dominant osteogenesis imperfecta initially described by Fisher *et al*. (30), exhibited a severe phenotype. The mutant zebrafish presented with high incidences of intervertebral ligament (IVL) mineralization, compressions, fusions, fractures and detachments. Additionally, the fin regeneration study showed a reduced fin length and altered fin shape after regeneration, indicating severely impaired collagen type I fiber formation and organization. Indeed, collagen type I is overmodified and retained in the endoplasmic reticulum (18). Furthermore, a delay in osteoblast differentiation and bone formation has been observed in caudal fin rays of the *col1a1a^dc124/+^* zebrafish (31), corroborating the results presented here. Histologically, the compression fracture phenotype of the *col1a1a^dc124/+^* mutant was noticeable and aligns with a recent study identifying vertebral compression fractures in the same model (32). As no clear fractures were observed in serial histological sections of this deformity, it appears that the bone of the vertebral centrum, i.e. the autocentra, buckles rather than fractures. This observation further corroborates that bone stiffness is reduced (17). This bone collapse should not occur without severe concurrent changes of the notochord. Whole mount-stained mutant zebrafish showed notochord mineralization, which presented histologically as fibrous tissue in the central part of the intervertebral discs extending into the notochord of the vertebrae, consisting with previous findings by Fiedler *et al*. (17) and Gioia *et al*. (18). This fibrous tissue has been shown to be a combination of collagen type I and keratins in a vertebral disc degeneration zebrafish model (33). The notochord septum was highly vacuolated and cartilage tissue was regularly present inside the notochord. Both the vacuoles of the chordocytes in the notochord septa and the cartilage provide support to tissues via hydrostatic pressure (34,35). Replacement of notochord by cartilage cells is a known disease mechanism in teleost fish (32,36). Combining previous findings with the results presented here on reduced bone stiffness and fragility, along with alterations in the notochord, it can be presumed that the tissue changes in the notochord function to stabilize the axial skeleton. Interestingly, the notochord changes observed in the current study were more severe compared to those reported by Cotti *et al*. (32). However, while Cotti and colleagues (32) used 3-month-old zebrafish, older zebrafish were used in the current study indicating that age may explain the differences in observed severity between the two studies. Alternatively, or in addition to age, the mineralization of the notochord core, as well as the stabilizing chordocytes and cartilaginous tissues, may result in a stiffer vertebral column. This rigidity may shift mechanical loads during swimming to other areas of the vertebral column, exacerbating deformities such as compressions and fractures, as observed in an osteoarthritis zebrafish model (33). One of the hallmark morphological deformities observed in whole mount stained chihuahua mutants was mineralization of the intervertebral ligament (IVL). Remarkably, the collagen present in the IVL as observed on TEM images was disorganized to such an extent that the mature collagen layer’s plywood-like structure was no longer recognizable. Additionally, a less compact packing of collagen fibers was evident. The increased number of chordoblasts and higher elastin grade in the mutant IVL further support these observations. Taken together, these findings indicate that both the notochord components (chordoblasts and elastin grade), and the collagen type I ligament are heavily altered in this mutant. The observed higher number of chordoblasts in the chihuahua mutant and the likely altered production of collagen type II in the notochord sheath may further explain a compensation mechanism aimed at stabilizing the vertebral column. The disorganization and especially the less condense packing of collagen type I fibers likely contribute to the ectopic mineralization of the IVL. Studies in salmon and zebrafish using different dietary phosphorous levels shave shown that collagen type I mineralization can be decoupled from collagen type I production and its processing into an unmineralized matrix (37,38). These experiments demonstrate that bone formation and mineralization can occur independently from each other. Importantly, less densely packed collagen type I fibers provide the space for mineral nucleation to take place (39). Taken together, the increased potential for mineralization initiation, along with semi-autonomous mineralization may explain the ectopic mineralization in the IVL, simply because the opportunity for mineralization is present at this location. Alternatively, this impaired mineralization may have a different, yet unknown mechanism.

The *col1a1a^dc124/+^* mutant, carries the same mutation identified in a human patient with severe and heavily deforming OI type III (40) and can be classified to the same OI type III, highlighting the utility of this zebrafish model for studying this specific form of OI. Although there is some variation in the phenotype of this zebrafish model, the phenotypic range is limited to a set of severe phenotypes that are very similar in appearance across all levels of morphological analyses. This observation, based on the results presented here, confirms the chihuahua mutant as an ideal model for OI treatment experiments as recently published (32,38).

Lastly, the *col1a2^mh15/+^* mutants exhibited notochord and IVL mineralization, vertebral detachments, fusions and scoliosis, consistent with findings by Gistelinck *et al* (23). Morphologically, the most striking feature of this mutant is the wide range of phenotypes, from moderate to very severe, reflected in the large variability of deformity scores, especially the total mineralization and ectopic ossification scores. The extensive ectopic mineralization in this mutant corresponds with the clinical manifestation of hyper-mineralization in patients with severe OI (41–43). Fin regeneration results also reflect the phenotypic variability in this mutant, showing morphological differences from moderate to very severe. Strikingly, the fin phenotype of the *col1a2^mh15/+^* severe mutants is even more severe compared to the already severe phenotype present in the *col1a1a^dc124/+^*mutants, indicating both collagen type I and cellular dynamic changes. Gistelinck and colleagues (23) noted slight or no collagen overmodification in this mutant, possibly due to the selection of a mixture of phenotypes or a bias towards less severe phenotypes for their analyses. Histologically, the phenotypes ranged from mildly altered vertebral endplates and notochord tissue to very severe alterations of the notochord tissue and IVL. Remarkably, in a very severe case, the detachment deformities, typically observed in vertebral arches, occurred at the level of an entire vertebra. Serial histology revealed the complete interruption of the vertebral column and the replacement of the notochord and bony tissue with fibrotic tissue high in collagen fibers. This fibrotic tissue likely stabilizes the vertebral column. Such a severe deformity provides further evidence of both collagen fiber alterations and cellular dynamic changes. Ultrastructural analysis revealed an increased number of cells in the IVL accompanied by disorganized collagen type I fibers with elevated elastin and autocentrum grades. Notably, TEM analyses indicated that the changes in cellular numbers are more pronounced than the alterations in collagen fiber organization. This finding further supports the hypothesis that the *col1a2^mh15/+^* is affected by both collagen fiber disorganization and altered cellular dynamics. In general, *col1a2^mh15/+^* mutants, documented in detail for the first time here, exhibit phenotypes on the more severe end of the spectrum, representing the most detrimental phenotype among the three analyzed mutants. The zebrafish mutant displays heavily distorted, misshapen, and improperly mineralized axial skeletons, characteristic of OI type II (23). This aligns with human data, where glycine substitutions in the homologous region of α2(I) are associated with lethal OI in human patients due to severe skeletal malformations early in fetal development, along with other extra-skeletal anomalies (40). Similar phenotypic variability as observed in this zebrafish mutant has been reported in human OI patients by Garibaldi *et al*. (2). In their study, 56.2% of 112 glycine residue substitutions for bulky amino acids (such as aspartic acid, often associated with lethal OI) in the type 1 collagen α-2 chain resulted in variable non-lethal type I-IV OI phenotypes. Such intra-familial variability is also seen with other genetic mutations in OI, such as those in *FKBP10*, which produce a range of phenotypes from mild to severe (7). Further exploration of the phenotypic variability in *col1a2^mh15/+^* mutants is needed. Researchers must investigate mild and severe phenotypes separately, which was not done here, possibly explaining the lack of significant differences in deformity scoring and ultrastructural analyses. The severe phenotype of *col1a2^mh15/+^*is worse than the chihuahua phenotype, suggesting that significant differences would have been found if phenotypes were separated. The wide range of phenotypes could help identify modifier genes influencing phenotype severity, as shown in a recent study where *bmpr1aa* was identified as a modifier gene in an *fkbp10*-deficient zebrafish model for OI and Bruck syndrome (24).

To identify molecular signatures (‘biomarkers’) associated with the severity of phenotypic outcomes in the three mutants, an MS-based proteome analysis was performed on the vertebral column of mutants versus wild-type siblings. In *col1a1a^mh13/+^*mutants, six proteins were upregulated, including three matrisome components: Col22a1, Serpine2, and Olfml3a. Col22a1 is involved in skeletal and cardiac muscle junctions and osteoblast regulation (44,45). Serpine2 is linked to cardiovascular diseases, fibrosis, and cancer (46). Olfml3a modulates BMP signaling and is associated with angiogenesis and tumor growth (47). In *col1a1a^dc124/+^* mutants, 13 differentially expressed proteins were identified, with 11 upregulated and two downregulated. Six of these are part of the zebrafish matrisome, including Fn1b, Col8a1a, Bcan, Prg4b, HSP47, and PEDF. Notably, *Serpinf1*, downregulated in these mutants, encodes PEDF, which regulates osteoclast activity and is linked to OI type VI (48,49). In *col1a2^mh15/+^*mutants, 61 proteins were upregulated and four downregulated. Several were linked to the zebrafish matrisome, including Bcan, Col8a1a, Ecm2, Vtna, Ihha, Lgals3b, LH2, HSP47, Shha, and Crtap. These proteins are involved in various functions such as cell proliferation, bone growth, inflammation, and collagen biosynthesis (50–53). Interestingly, Bcan, HSP47, and Col8a1a were upregulated in both *col1a1a^dc124/+^*and *col1a2^mh15/+^* mutants but not in the mildly affected *col1a1a^mh13/+^*mutants, suggesting their potential as biomarkers for severe OI. The *SERPINH1* gene encodes heat shock protein 47 (HSP47), a collagen-specific chaperone that stabilizes folded procollagen and facilitates its transfer to the Golgi apparatus (54). Mutations in *SERPINH1* cause OI type X, a severe form often associated with lethal outcomes (55). In zebrafish, *serpinh1b* supports fin regeneration by mediating Cx43-dependent skeletal growth. Upregulated Hsp47b expression has been observed in chihuahua zebrafish at 5 days post fertilization, localized to the tail fin fold, skin, and intersomitic space (18). Increased expression of *serpinh1b* was also found in fin tissue of severely, but not in mildly affected *fkbp10^-/-^* zebrafish mutants (24). Elevated HSP47 levels, along with increased colocalization with mutant type I collagen, were observed in OI patient cells and murine models (56,57). This upregulation may represent an adaptive effort to ensure proper collagen synthesis or reflect impaired movement of mutant collagen (18). Gene ontology analysis highlights HSP47’s role in collagen fibril organization. Both *col1a1a^dc124/+^*and *col1a2^mh15/+^* mutants, but not *col1a1a^mh13/+^*mutants, show disorganized collagen fibrils, suggesting HSP47’s contribution to phenotype severity. BCAN or brevican is involved in brain ECM formation. *Bcan* knock-out mice exhibit brain abnormalities, including impaired long-term potentiation (58). In dogs, a bcan microdeletion is linked to episodic flailing syndrome (59). In Xenopus, *brevican* expression in notochord buds suggests its role in notochord development (60,61). In zebrafish, Brevican may also play a role in correct notochord development, which we showed was severely affected in both the *col1a1a^dc124/+^* and *col1a2^mh15/+^* mutants, and its maintenance, although this remains to be investigated. Type VIII collagen, encoded by *col8a1*, is a key component of the Descemet’s membrane of corneal endothelial cells (62). Mutations in *COL8A1* are associated with eye abnormalities in OI, including decreased cornea thickness (*62*). Upregulation of Col8a1 in severe OI models suggest its role in skeletal phenotypes (63). Zebrafish lacking *col8a1a* exhibit vertebral malformations, indicating its involvement in ECM structure and function (64). The upregulation of Bcan, HSP47, and Col8a1a in severe *col1a1a^dc124/+^* and *col1a2^mh15/+^*mutants indicates their potential as biomarkers for severe dominant OI, contributing to diagnostics, clinical follow-up, and treatment guidance.

While the findings in this study are both new and interesting, it is important to acknowledge certain limitations when interpreting the results. The shotgun mass spectrometry, while valuable, has inherent limitations, such as difficulty in detecting low-abundance proteins and identifying poorly annotated ones. As a results, only a subset of the proteins present in the vertebral column was detected, limiting the ability to fully analyze molecular mechanisms underlying disease severity. Developing a deformity scoring matrix tailored to specific deformities observed in zebrafish models for OI and other connective tissue diseases is a significant strength. Although the scoring results presented here focused on disease-relevant phenotypes, the matrix can be expanded to include more naturally occurring deformities. While using only three samples per genotype per mutant model for serial histology might seem low and could be perceived as a limitation, it is actually a strength of this research. Differential interference contrast (DIC) light on unstained paraffin sections allowed for the specific selection of histological slides to be stained with different stains, validating observations within the same tissue. The number of TEM samples, six per genotype per mutant model, is notably high for this time-consuming expert technique. This detailed, intensive approach enabled the authors to connect morphological phenotypic data to histological and ultrastructural data, thus understanding each phenotype and deformity in greater detail.

Despite these limitations, this study opens avenues for future research. The identification of potential biomarkers for phenotypic severity in dominant OI requires further investigation to validate their relevance and role in the studied condition, particularly in human cell tissue samples such as blood serum, bone tissue or bone cell cultures. Their roles in ECM organization, collagen processing, and skeletal development further support their potential involvement in the pathogenesis of severe OI phenotypes. However, further research is needed to fully elucidate the molecular pathways and mechanisms by which these differentially expressed proteins influence the severity and variability of the mutant phenotypes in the different OI models.

In conclusion, our comprehensive phenotypic, morphological, (ultra)structural, and molecular analyses of three distinct dominant OI zebrafish models with different glycine substitutions in the collagen type I genes has provided valuable insights into the phenotypic variability associated with this condition. A detailed phenotypic scoring matrix for disease models, which not only serves as a robust tool for assessing skeletal deformities in OI zebrafish models but also holds potential for evaluating other skeletal diseases affecting the vertebral column has been successfully established. The findings presented here indicate that increasing skeletal severity in dominant OI correlates with a higher incidence of skeletal deformities and abnormalities. This, in turn, can be associated with thinner bones and increased disorganization of collagen fibrils, notochord alterations and ectopic mineralization, elastin deposit changes, and cellular dynamic changes in the notochord and IVL. These insights, combined with genetic information, have the potential to predict phenotypic outcomes in dominant forms of OI. This study further reports a remarkable intra-familial phenotypic variability in one of the mutants which holds potential for future approaches that could help in understanding the underlying mechanisms of this variability and the identification of modifier genes. Finally, this research led to the identification of three potential molecular markers, HSP47, Col8a1 and Bcan, that could serve as indicators of disease severity. Such molecular markers possess diagnostic value and also offer predictive insights into the progression of clinical presentations and play a crucial role in guiding treatment. However, further investigation is required to validate their clinical relevance. Collectively, the present findings contribute to a deeper understanding of the variable presentation of dominant forms of OI and paves the way for future studies aimed at improving diagnostic and therapeutic approaches.

## Materials and Methods

### Zebrafish husbandry and maintenance

The *col1a1a^mh13/+^* (G1093R; a1-Gly931Arg, p.Gly1093Arg), *col1a2^mh15/+^* (G882D; a-2Gly802Asp, p.Gly882Asp) and the *col1a1a^dc124/+^* (G574D; a1-Gly574Asp, p.Gly736Asp) mutants were generated in a forward genetics screen (30,65) and described in Gistelink *et al*, (23). All zebrafish were kept in the Core Zebrafish Facility Ghent (ZFG) and housed in a Zebtec semi-closed recirculation housing system provided with reverse osmosis purified water at a constant temperature (27°C), pH (7.5), conductivity (550μS) with a light(14h)/dark(10h) cycle. They were fed with Zebrafeed (Sparos) and Gemma Micro dry food (Inve) in the morning and Micro Artemia (Ocean Nutrition) in the afternoon. All animal experiments were performed according to EU Directive 2010/63/EU, permit no. ECD 23/27 (Ghent University). Pain, distress, and discomfort were minimized as much as possible. Numbers of specimens that were selected for each analysis are listed in Table 1.

**Table 1.**
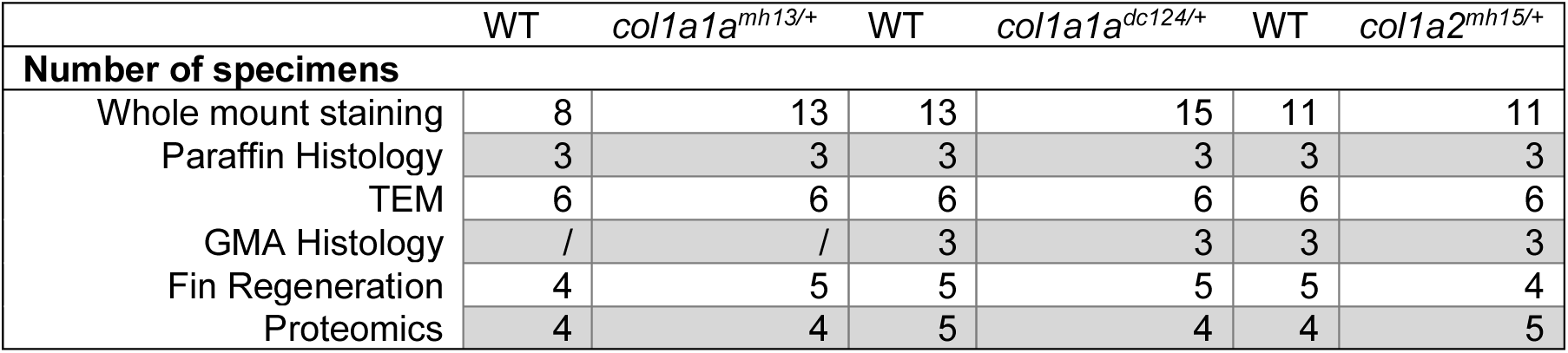
Number of adult zebrafish used in the different analyses. WT = wild type siblings.

|  | WT | *col1a1a<sup>mh13/+</sup>* | WT | *col1a1a<sup>dc124/+</sup>* | WT | *col1a2<sup>mh15/+</sup>* |
| --- | --- | --- | --- | --- | --- | --- |
| **Number of specimens** |  |  |  |  |  |  |
| Whole mount staining | 8 | 13 | 13 | 15 | 11 | 11 |
| Paraffin Histology | 3 | 3 | 3 | 3 | 3 | 3 |
| TEM | 6 | 6 | 6 | 6 | 6 | 6 |
| GMA Histology | / | / | 3 | 3 | 3 | 3 |
| Fin Regeneration | 4 | 5 | 5 | 5 | 5 | 4 |
| Proteomics | 4 | 4 | 5 | 4 | 4 | 5 |

**Table 2.** Measured parameters during TEM live imaging and on TEM micrographs compared between mutant and WT. All parameters are explained in Table S2. The indication of ‘=’ means the average values were ± 10% between mutants and WT, ‘>’ and ‘<’ means that the average value was ± 10-25% between mutants and WT, ‘>>’ and ‘<<’ means that the average value was ± 25-40% between mutants and WT and ‘>>>’ and ‘<<<’ means that the average value was ± more than 40% between mutants and WT. * indicates significant differences.

| Class | Parameter | Structure | *col1a1a<sup>mh13/+</sup>* | *col1a1a<sup>dc124/+</sup>* | *col1a2<sup>mh15/+</sup>* |
| --- | --- | --- | --- | --- | --- |
| Class 1:<br>number of<br>cells | Chordoblasts | Notochord | = | >>> | >>> |
|  | Cells intervertebral ligament | Ligament | >> | = | >>> |
|  | Cells on mature collagen | Ligament | >> | << | >> |
|  | Cells on bone | Bone | > | > | = |
| Class 2:<br>thickness of<br>collagen<br>layers | Mature collagen thickness | Ligament | = | <<< | << |
|  | Number of layers | Ligament | >> | << | < |
|  | Longitudinal layers | Ligament | = | << | > |
|  | Circumferential layers | Ligament | << | << | > |
|  | Bone thickness | Bone | < | <<<*** | << |
| Class 3:<br>grade of key<br>structures | Angle | Notochord | >> | = | > |
|  | Elastin grade | Notochord | >> | >> | >> |
|  | ER grade | Ligament | >> | >> | >> |
|  | Autocentrum grade | Bone | >> | >>>* | >>> |

### Whole mount Alizarin Red S staining

Adult mutant zebrafish and wild-type siblings were stained using the Alizarin red S for mineralized tissue following a protocol established by Sakata-Hage *et al*. (66) and adjusted as described by Bek *et al.* (67)(SI text). Images of stained zebrafish were taken using a Leica M165FC binocular stereomicroscope with the LAS V4.3 software (Leica Microsystems). Following consistent dissection, the underlying skeleton was exposed. Lateral observation was employed for most skeletal assessments, except for spinal curvature evaluation, where the ventral side was used. Brightfield and fluorescent modes, using the DSR DsRed Dichroic Mirror filter, were used to examine each specimen.

### Phenotypic scoring matrix

A novel scoring matrix, designed for reproducible skeletal phenotyping, was developed based on an offspring of fifty, five-months old *col1a2^mh15/+^*zebrafish generated from a single parent couple. Within this family, a total of 39 unique deformities were identified in the vertebral column, comprising both natural (Figure S1) and pathogenic variations (Figure S2)(68). The final scoring matrix considered only the pathological deformities (Table S1) and the absence (value 0) or presence (value 1) of these deformities in each vertebral body was noted, excluding the first four vertebrae comprising the Weberian apparatus, and the last four vertebrae comprising the preural and ural vertebrae.

This scoring matrix was used to compare three mutant families, *col1a1a^mh13/+^*, *col1a1a^dc124/+^*, and *col1a2^mh15/+^*, to their respective wild-type counterparts, aged 8 to 11 mpf, using Alizarin Red S staining. Deformity frequency was calculated as a percentage of deformed vertebrae, with data combined to determine the overall deformity score. Statistical comparisons of phenotypic, total mineralization score, and ectopic ossification scores were performed using One-Way ANOVA with Tukey’s post hoc analysis in GraphPad Prism version 9.4.0. (www.graphpad.com). High-magnification images of representative specimens were captured using an Axiozoom V16 stereomicroscope (Carl Zeiss, Oberkohn, Germany) and Zen Pro software, with composite images generated using Autostitch demo version 2.3 (69).

### Fin regeneration

To evaluate fin regeneration, all mutants were outcrossed with a transgenic *osx:*Kaede fluorescent line, specifically labeling osteoblasts. The caudal fin was amputated in adult zebrafish under general anesthesia (160 mg/L MS-222). Photoconversion of native green Kaede fluorescence in fins at 0 days post-amputation (dpa) was performed using a Zeiss Observer microscope with 405 nm UV exposure (DAPI) for 30 seconds. Imaging of the lateral side of the fins at 8- and 21-days post-amputation (dpa) was performed using GFP3 and dsRed filters on a Leica M165FC stereomicroscope, equipped with a DFC450C camera and controlled by LAS V4.3 software (Leica Microsystems). Image processing was carried out using Fiji software (version 2.14.0).

### Histological analysis

#### Paraffin histology

Embedding and analysis was performed following similar methodology as described in Gistelinck *et al*. (23)(SI text)

#### GMA histology

Specimens at 11 mpf used for histological analysis were first euthanized by tricaine overdose and fixed for 24 hours in 2.5% PFA, 0.1 M sodium cacodylate buffer (pH 7.4) and 0.001% CaCl2 at 4°C. Bone mineral detection was carried out on histological sections obtained from non-decalcified samples embedded in glycol methacrylate (GMA), according to Witten *et al*. (70)(SI text). All sections were observed and photographed using a Axio Imager-ZI microscope (Carl Zeiss, Oberkohn, Germany) equipped with a Axiocam 503 Color camera (Carl Zeiss, Oberkohn, Germany) and Zen Pro 2 software (Carl Zeiss, Oberkohn, Germany).

#### Transmission electron microscopy

Specimens at 7 mpf were fixed, decalcified and processed for TEM (SI text). Several parameters of the intervertebral ligament and end plates were measured. These parameters were grouped into three main categories namely (i) the number of cells, (ii) the thickness and number of collagen layers and (iii) the grading of key structures (Table S2). An average value was calculated for both mutant and wild type siblings, respectively. A One-way ANOVA was used to compare the mutants and their matched wild-types in GraphPad Prism version 9.4.0 (GraphPad Software, San Diego, California, USA, www.graphpad.com). Raw TEM data can be found in Dataset S2. To construct table 2, the averages between mutant and wild type siblings were compared. The largest differences were marked with triple symbols (<<< or >>>>), moderate with double (<< or >>), smallest with single (< or >), and equal values with (=).

### Mass spectrometry

#### Sample preparation

Adult zebrafish and respective sibling controls were euthanized using a 25x tricaine solution and vertebral bone was dissected. Excess soft tissue was removed after incubation in Accumax solution (Sigma-Aldrich) for 2 hours. Proteins were extracted using Trizol (71)(SI text).

#### LC-MS/MS analysis

Peptides were re-dissolved in 20 µl loading solvent A (0.1% TFA in water:ACN (98:2, v:v)) moments before analysis. The LC-MS/MS analysis protocol is described in the SI text.

#### Data analysis

Analysis of the mass spectrometry data was performed using MaxQuant (version 2.0.3.0) with default search settings which included a false discovery rate set at 1% at the PSM and protein level. Spectra were searched against the *Danio rerio* reference proteome (version of February 2021, UP000000437). A more detailed description of the data analysis is provided in the SI text. The mass spectrometry proteomics data have been deposited to the ProteomeXchange Consortium via the PRIDE partner (72) repository with the dataset identifier PXD058825.

Gene Ontology (GO) enrichment analysis was performed using the clusterProfiler R package (R version 4.4.1) to identify over-represented GO terms in the dataset. Ensembl gene IDs were mapped to Entrez gene IDs using the BioMart database (biomaRt package) with the *Danio rerio* gene dataset. The analysis was conducted for each fish group separately. GO enrichment was carried out using the enrichGO function and GO processes with an adjusted p-value (FDR) ≤ 0.05 were considered significant.

Protein-protein interaction (PPI) networks were constructed using STRINGdb (v11.0) for *Danio rerio* (species ID: 7955) with a confidence score threshold of 400, integrating both physical and functional interactions. The resulting network was visualized using the igraph package, with node colors indicating gene regulation direction and experimental groups.

## Acknowledgments

LS acknowledges funding from the Geconcerteerde onderzoeksacties funding from Ghent University BFO.GOA.20210004.07. SD acknowledges funding from the fonds wetenschappelijk onderzoek (FWO) FWO.OPR.2020.0023.01 and the Care 4 Brittle Bones foundation. We would like to thank the Zebrafish Facility Ghent (ZFG) Core and TEM Core Facility at Ghent University.

