## Supplementary Material for "Decoding Phenotypic Variability in Osteogenesis Imperfecta: Zebrafish as a Model for Molecular and Ultrastructural Insights"

**This PDF file includes:**

Supporting text
SI References
Figures S1 to S2
Tables S1 to S2
Legends for Datasets S1 to S4

**Other supporting materials for this manuscript include the following:**

Datasets S1 to S4

### Supporting Information Text

### Extended Materials and Methods

#### Alizarin red S staining protocol

The euthanized and fin-clipped specimens were immersed in a fixative solution (4% formaldehyde, 5% polyoxyethylene-(10)-octylphenyl ether [Triton X-100] and 1% KOH) for 10 days at room temperature. After this, the specimens were immersed in a clearing solution (20% ethylene glycol, 5% Triton X-100 and 1% KOH) for 24 hours, followed by bone-staining medium (20% ethylene glycol and 1% KOH) for 30 minutes and thereafter bone staining solution (0.05% alizarin red S [A3757, Sigma-Aldrich, Darmstadt, Germany], 20% ethylene glycol and 1% KOH) for 24 hours. Next, the specimens were rinsed with distilled water after which the scales of the zebrafish were removed using a cotton swab. To remove background staining, the specimens were washed in a de-staining solution (20% polyoxymethylene (20) sorbitan monolaurate [Tween 20] and 1% KOH) for 48 hours. Subsequently, the specimens were moved through a graded series of glycerol (20% to 100%) and stored in 100% glycerol.

#### Paraffin histology

Adult zebrafish (n=3 per mutant and respective sibling wild-type) were fixed in modified Davidson's Fixative for 24 hours and then 10% neutral buffered formalin overnight. Thereafter, the specimens were decalcified in citric acid (45% formic acid, 20% sodium citrate and water, 1:1:2) for four to six hours at room temperature. The specimens were subsequently dehydrated in series of ethanol (70% overnight, 90% two hours, 99% overnight) and transferred to xylene for eight hours. The samples were then impregnated with paraffin at 65°C overnight. The whole specimens were embedded in paraffin using a Leica Embedding machine. Parasagittal serial sections were cut at 4 µm using a Thermo Scientific microtome. Sections were stained with Meyer's acid Hematoxylin and Eosin (Sigma Aldrich) for assessment of general structures, Masson's trichrome (Sigma Aldrich) staining collagen type I deep red, Verhoeff-Van Gieson staining elastic fibers black, collagen type I red and cartilage collagen type II pink (NovaUltra) and Heidenhain's AZAN trichrome (Morphisto, Germany) staining collagen type I deep blue (and red to varying degree), elastin vibrant red and cartilage collagen type II light blue, following the protocols outlined by the manufacturer.

#### GMA histology

Briefly, the region of interest (last 2 abdominal and first three caudal vertebrae) was dissected from the whole specimens and dehydrated in a graded series of acetone (30%, 50%, 70%, 90%, 100%) in 45-minute increments. Samples were then impregnated with GMA monomer solution (80 mL (2-hydroxyethyl)-methacrylate, 12 mL ethylene glycol monobuthyl ether, 270 mg benzoyl peroxide) for one hour. For the second step of impregnation a fresh monomer solution was used for one week. For embedding 2% catalyst (1 mL N,N-dimethylaniline, 10 mL poly-ethylenglycole-200) was added to the monomer solution. Specimens were then embedded in a mold. Polymerization took place at 4°C for 24 hours and was completed within another 48 hours at room temperature. Four µm sagittal sections were cut on a Microm HM 360 (Marshall Scientific, Hampton, NH, USA) automated microtome and were stained using the Von Kossa/Van Gieson staining protocol for phosphate detection: 1% $AgNO_3$ (45 min under UV light); $dH_2O$ (10 min, twice); 3% $Na_2S_2O_3$ (5 min); $dH_2O$ (10 min, twice); Van Gieson counterstain (5 min); $dH_2O$; air-drying and DPX mounting (1). Additionally, serial sections were stained with a 0.5% aqueous solution of Alizarin red S for 4 minutes to demonstrate the presence of calcium in the bone.

#### Transmission electron microscopy

Specimens were fixed for 24 hours in 2.5% PFA, 1.5% glutaraldehyde, 0.1 M sodium cacodylate buffer (pH 7.4) and 0.001% $CaCl_2$ at 4°C. Fixed fish were decalcified in 0.1 M EDTA for 14 days. The decalcification solution was changed every 3 days. Specimens were subsequently rinsed in 0.1 M sodium cacodylate buffer with 10% saccharose and then post-fixed for 2 hours in 1% $OsO_4$ solution in 0.1 M cacodylate buffer containing 3% saccharose. After rinsing in buffer, samples were dehydrated in a series of graded ethanol solutions and embedded in epon epoxide medium. Semithin 1 µm sections were cut on a Microm HM360 microtome (Marshall Scientific, Hampton, NH, USA), stained with toluidine blue at pH 9 for 2 min (0.5% toluidine blue, 1% Na2B4O7 in dH2O), rinsed with H2O and mounted with DPX. For transmission electron microscopy (TEM) analysis, ultrathin sections (about 80 nm) of the region of interest were prepared on an UltracutE ultramicrotome (Reichert-Jung, Buffalo, NY, USA), contrasted with uranyl acetate and lead citrate (2) and analyzed with a Jeol JEM 1010 transmission electron microscope (Jeol Ltd., Tokyo, Japan) operating at 60 kV. Microphotographs were taken with a Veleta camera (Emsis, Muenster, Germany).

**Mass spectrometry – sample preparation**

The tissue was transferred to 1 ml Trizol, homogenized using the TissueLyser II (Qiagen) and incubated on ice for 5 min. After centrifugation, the supernatants was collected. Chloroform was added and the samples were again incubated on ice for 5 min before another round of centrifugation. The upper aqueous RNA phase was separated, and DNA was precipitated using ethanol. Following centrifugation, the supernatants was collected, and proteins were precipitated using isopropanol. The samples were centrifuged, resulting in the formation of a protein pellet. The protein pellet was washed with 0.3 M guanidine hydrochloride solution and ethanol twice, after which the pellet was air-dried. A lysis buffer containing 10% sodium dodecyl sulfate (SDS) and 100 mM triethylammonium bicarbonate (TEAB; pH 8.5) was added to the samples to a final concentration of 4.5% SDS and 50 mM TEAB. The samples were sonicated for 15 minutes and proteins were reduced with 15 mM dithiothreitol and incubated for 30 minutes at 55°C and then alkylated with 30 mM iodoacetamide and incubated for 15 minutes at RT in the dark. Phosphoric acid was added to a final concentration of 2.75% and subsequently samples were diluted 7-fold with a binding buffer (90% methanol in 100 mM TEAB, pH 7.55). After loading 100 µg protein to S-trap micro-columns (Protifi) using centrifugation for 30 s at 4.000 g, the columns were washed three times with 150 µl binding buffer, after which trypsin (1/100, w/w) was added for digestion overnight at 37°C. The next day, peptides were eluted three times, first with 40 µl 50 mM TEAB, then with 40 µl 0.2% formic acid (FA) in water and finally with 40 µl 0.2% FA in water/acetonitrile (ACN) (50/50, v/v). Eluted peptides were dried completely by vacuum centrifugation.

**Mass spectrometry – LC-MS/MS analysis**

From each sample, 1 µl of the sample was injected for LC-MS/MS analysis using an Ultimate 3000 RSLCnano system in-line, connected to a Q Exactive HF BioPharma mass spectrometer (Thermo). Trapping was performed at 10 µl/min for 4 min in loading solvent A on a 20 mm trapping column (made in-house, 100 µm internal diameter [I.D.], 5 µm beads, C18 Reprosil-HD, Dr. Maisch, Germany). The peptides were separated on a 200 cm µPAC™ column (C18-endcapped functionality, 300 µm wide channels, 5 µm porous-shell pillars, inter pillar distance of 2.5 µm and a depth of 20 µm; Thermo) at a constant temperature of 50°C. Peptides were eluted by a linear gradient reaching 55% MS solvent B (0.1% FA in water/acetonitrile (2:8, v/v)) after 100 min and 70% MS solvent B at 125 min, followed by a 5-minute wash at 70% MS solvent B and re-equilibration with MS solvent A (0.1% FA in water). The first 15 min the flow rate was set to 500 nl/min after which it was kept constant at 300 nl/min. The mass spectrometer was operated in data-dependent mode, automatically switching between MS and MS/MS acquisition for the 16 most abundant ion peaks per MS spectrum. Full-scan MS spectra (375-1500 m/z) were acquired at a resolution of 60.000 in the Orbitrap analyzer after accumulation to a target value of 3,000,000. The 16 most intense ions above a threshold value of 13.000 were isolated with a width of 1.5 m/z for fragmentation at a normalized collision energy of 28% after filling the trap at a target value of 100.000 for maximum 80 ms. MS/MS spectra (200-2000 m/z) were acquired at a resolution of 15.000 in the Orbitrap analyser. The polydimethylcyclosiloxane background ion at 445.120028 Da was used for internal calibration (lock mass) and QCloud was used to control instrument longitudinal performance during the project (3,4).

**Mass spectrometry – Data analysis**

During the main search, the mass tolerance for precursor and fragment ions was set to 4.5 and 20 ppm, respectively. Enzyme specificity was set as C-terminal to arginine and lysine, allowing cleavage at proline bonds with a maximum of two missed cleavages. Variable modifications were set to oxidation of methionine residues and acetylation of protein N-termini. Matching between runs was enabled with a matching time window of 0.7 minutes and an alignment time window of 20 minutes. Only proteins with at least one unique or razor peptide were retained. Proteins were quantified by the MaxLFQ algorithm integrated in the MaxQuant software. A minimum ratio count of two unique or razor peptides was required for quantification. In total, 26,274 unique peptides were identified corresponding to 3.346 proteins (Dataset S3). Further data analysis was performed with an in-house R script, using the proteinGroups table from MaxQuant as input. Reverse database hits were removed, LFQ intensities were log2 transformed and replicate samples were grouped. Proteins with less than three valid values in at least one group were removed and missing values were imputed from a normal distribution centered around the detection limit (package DEP) leading to a list of 2.070 quantified proteins (Dataset S4-5) (5). To compare protein abundancies between pairs of sample groups (WT sibling of col1a1amh13/+ versus col1a1amh13/+, WT sibling of col1a1adc124/+ versus col1a1adc124/+ and WT sibling of col1a2mh15/+ versus col1a2mh15/+) statistical testing for differences between two group means was performed, using the package limma (6). Statistical significance for differential regulation was set to a false discovery rate (FDR) of <0.05 and fold change of >2.83- or <0.35-fold (|log2FC| = 1.5).

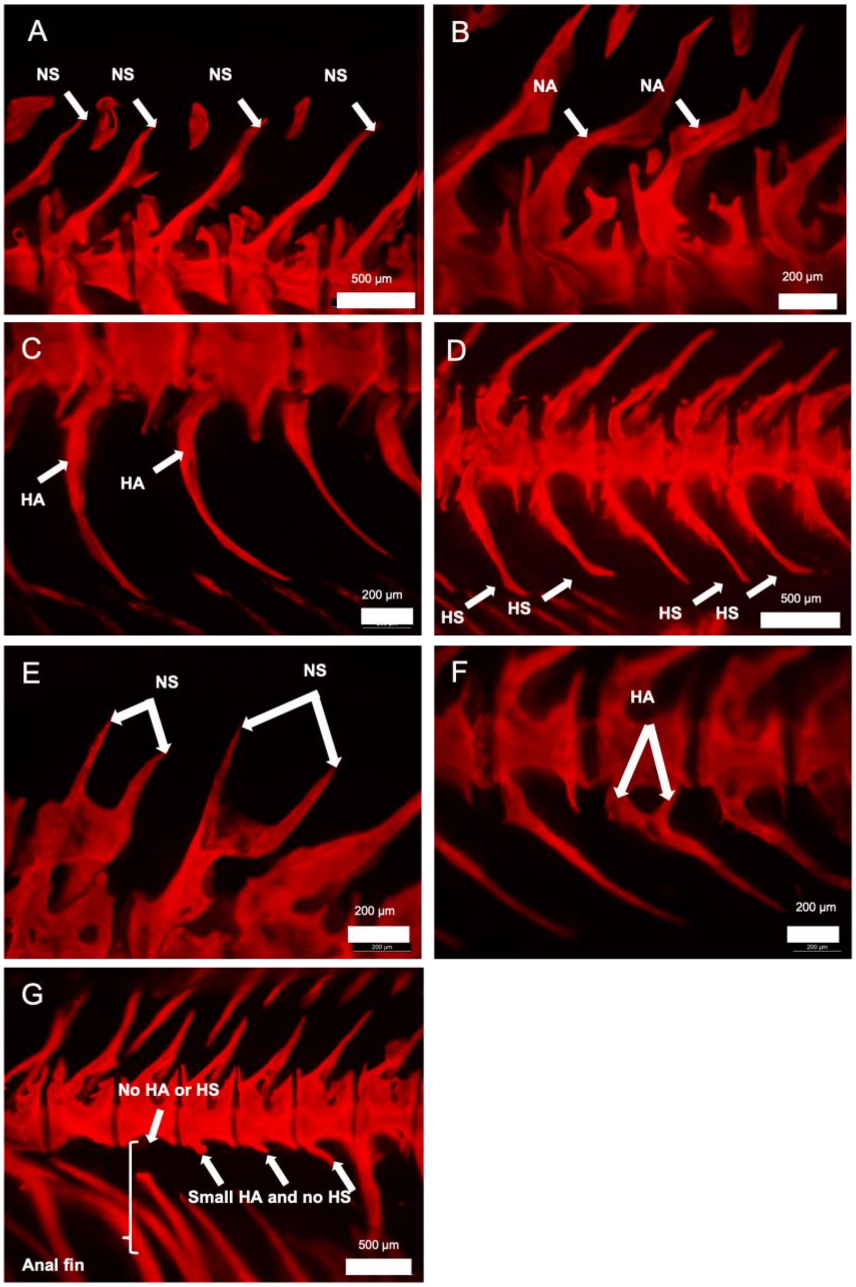


**Figure S1:** Natural variations in the zebrafish vertebral column. (A) bent neural spine. White arrows indicate a curve in the neural spines; (B) bent neural arch. White arrows indicate a curve in two neural arches; (C) bent haemal arch. White arrows indicate a curve in two haemal arches; (D) bent haemal spine. White arrows indicate a curve in four haemal spines. (E) double neural spine. White arrows indicate two double neural spines; (F) double haemal arch. White arrows indicate a double haemal arch originating from the same vertebral body; (G) lacking haemal arch and/or haemal spine. White bracket indicates anal fin rays while the most left white arrow indicates the lack of both a haemal arch and a haemal spine. The most right white arrows indicate three small haemal arches with no haemal spines attached to.

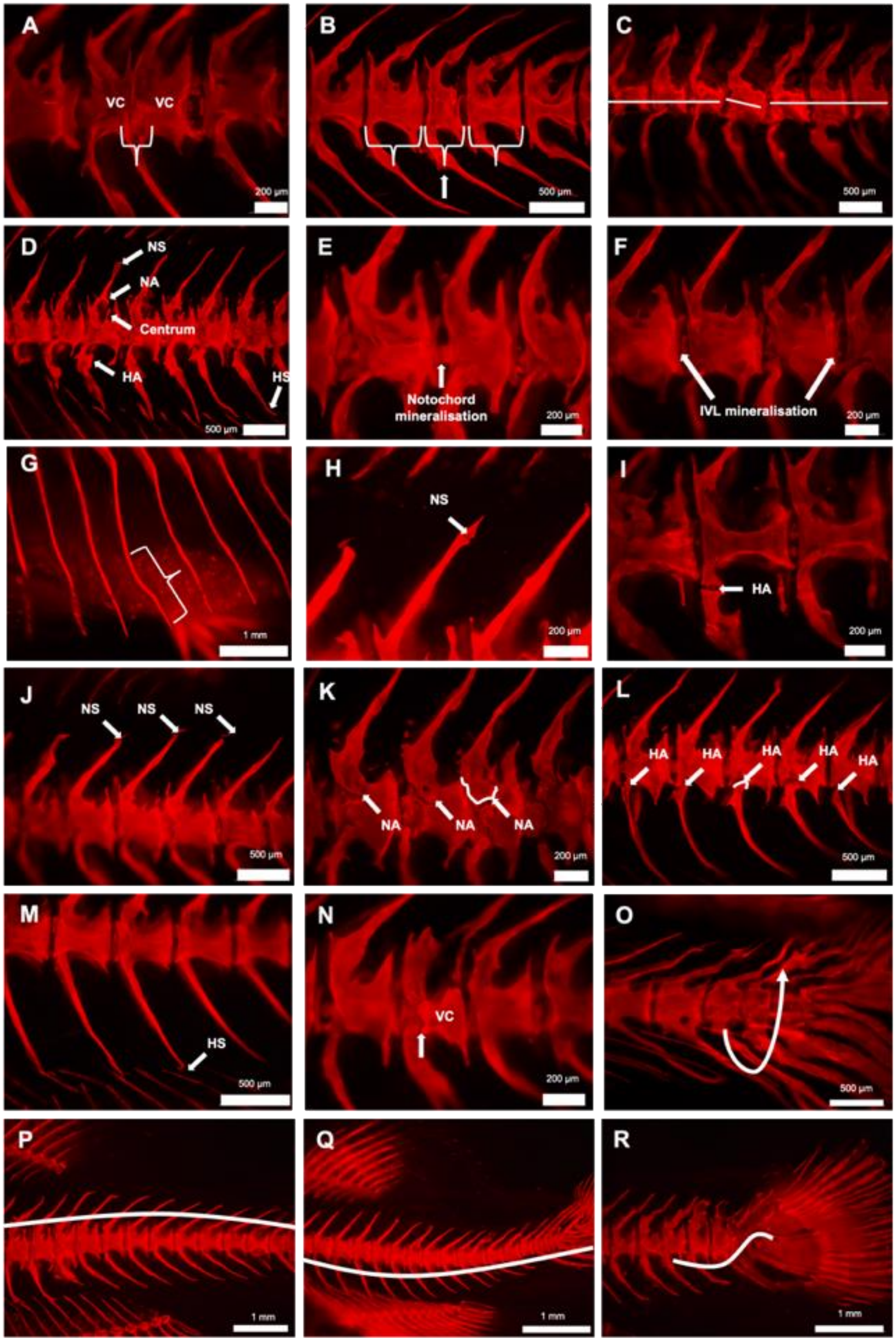
A
VC
VC
200 µm
B
500 µm
C
500 µm
D
NS
NA
Centrum
HA
HS
500 µm
E
Notochord mineralisation
200 µm
F
IVL mineralisation
200 µm
G
1 mm
H
NS
200 µm
I
HA
200 µm
J
NS
NS
NS
500 µm
K
NA
NA
NA
200 µm
L
HA
HA
HA
HA
HA
500 µm
M
HS
500 µm
N
VC
200 µm
O
500 µm
P
1 mm
Q
1 mm
R
1 mm

**Figure S2:** Pathological variations in the zebrafish vertebral column. (A) fusion of two vertebral centra. White bracket indicates the fusion and lack of intervertebral space; (B) compression of a vertebral centrum. White brackets indicate a compressed vertebral centrum in between normal sized vertebral centra; (C) vertical shift. White lines indicate a downward shift of one vertebral body, while the other vertebrae form a straight line. (D) ectopic ossification. White arrows indicate extra intramembranous ossification on the neural spine (NS), neural arch (NA), vertebral centrum (Centrum), haemal arch (HA) and haemal spine (HS); € mineralisation of internal notochord tissue. White arrow indicates staining of internal notochord tissue mineralisation within the intervertebral space (IVC); (F) intervertebral ligament mineralisation. White arrows indicate mineralisation of the notochord sheath and IVL tissue which can be seen on the surface of the intervertebral space. (G) curly rib. White bracket indicates a bend in the rib (H) fracture in neural spine. White arrow indicates callus formation in a neural spine which is indicative for a fracture; (I) fracture in haemal arch. White arrow indicates remodelled fracture lines in a haemal arch; (J) detachment of neural spine. White arrows indicate the detachment of three neural spines from their neural arches; (K) detachment of neural arch. White arrows indicate the detachment of three neural arches from their vertebral centra. White line indicates the base of the detached neural arch; (L) detachment of haemal arch. White arrows indicate the detachment of five haemal arches from their vertebral centra. White line indicates the base of the detached haemal arch; (M) detachment of haemal spine. White arrow indicates the detachment of a haemal spines from its haemal arch; (N) detachment of vertebral centrum. White arrow shows the fragmentation of the vertebral centrum; (O) torsion. White arrow indicates a rotation of the spine around the central axis causing the haemal arches and spines to be more superficially located than the neural arches and spines; (P) kyphosis. White line indicates a dorsal curvature of the spine; (Q) lordosis. White line indicates a ventral curvature of the spine; (R) scoliosis. White line indicates a lateral curvature of the spine.

**Table S1:** Phenotypic scoring: template used to establish the phenotypic scores. Colors define the part of the vertebral column (blue: precaudal vertebrae, green: caudal vertebrae, orange: caudal fin vertebrae, grey: caudal fin). All observable deformities are listed and scored (0=absent, 1=present).

| Animal | genotype | deformities | | | | | | | | | | | | | | | | | | | | | | | | | | | | | | |
|---|---|---|---|---|---|---|---|---|---|---|---|---|---|---|---|---|---|---|---|---|---|---|---|---|---|---|---|---|---|---|---|---|
| | | | 5 | 6 | 7 | 8 | 9 | 10 | 11 | 12 | 13 | 14 | 15 | 16 | 17 | 18 | 19 | 20 | 21 | 22 | 23 | 24 | 25 | 26 | 27 | 28 | 29 | 30 | 31 | 32 | 33 | 34 | 35 |
| | | total vertebrae | 1 | 1 | 1 | 1 | 1 | 1 | 1 | 1 | 1 | 1 | 1 | 1 | 1 | 1 | 1 | 1 | 1 | 1 | 1 | 1 | 1 | 1 | 1 | 1 | 1 | 1 | 1 | 1 | 1 | 1 | 0 |
| | | transitional | 0 | 0 | 0 | 0 | 0 | 0 | 0 | 0 | 0 | 0 | 0 | 0 | 0 | 0 | 0 | 0 | 0 | 0 | 0 | 0 | 0 | 0 | 0 | 0 | 0 | 0 | 0 | 0 | 0 | 0 | 0 |
| | | fusion | 0 | 0 | 0 | 0 | 0 | 0 | 0 | 0 | 0 | 0 | 0 | 0 | 0 | 0 | 0 | 0 | 0 | 0 | 0 | 0 | 0 | 0 | 0 | 0 | 0 | 0 | 0 | 0 | 0 | 0 | 0 |
| | | compression | 0 | 0 | 0 | 0 | 0 | 0 | 0 | 0 | 0 | 0 | 0 | 0 | 0 | 0 | 0 | 0 | 0 | 0 | 0 | 0 | 0 | 0 | 0 | 0 | 0 | 0 | 0 | 0 | 0 | 0 | 0 |
| | | vertical shift | 0 | 0 | 0 | 0 | 0 | 0 | 0 | 0 | 0 | 0 | 0 | 0 | 0 | 0 | 0 | 0 | 0 | 0 | 0 | 0 | 0 | 0 | 0 | 0 | 0 | 0 | 0 | 0 | 0 | 0 | 0 |
| | | curly ribs | 0 | 0 | 0 | 0 | 0 | 0 | 0 | 0 | 0 | 0 | 0 | 0 | 0 | 0 | 0 | 0 | 0 | 0 | 0 | 0 | 0 | 0 | 0 | 0 | 0 | 0 | 0 | 0 | 0 | 0 | 0 |
| | | detachment NS | 0 | 0 | 0 | 0 | 0 | 0 | 0 | 0 | 0 | 0 | 0 | 0 | 0 | 0 | 0 | 0 | 0 | 0 | 0 | 0 | 0 | 0 | 0 | 0 | 0 | 0 | 0 | 0 | 0 | 0 | 0 |
| | | detachment NA | 0 | 0 | 0 | 0 | 0 | 0 | 0 | 0 | 0 | 0 | 0 | 0 | 0 | 0 | 0 | 0 | 0 | 0 | 0 | 0 | 0 | 0 | 0 | 0 | 0 | 0 | 0 | 0 | 0 | 0 | 0 |
| | | detachment centrum | 0 | 0 | 0 | 0 | 0 | 0 | 0 | 0 | 0 | 0 | 0 | 0 | 0 | 0 | 0 | 0 | 0 | 0 | 0 | 0 | 0 | 0 | 0 | 0 | 0 | 0 | 0 | 0 | 0 | 0 | 0 |
| | | detachment HA | 0 | 0 | 0 | 0 | 0 | 0 | 0 | 0 | 0 | 0 | 0 | 0 | 0 | 0 | 0 | 0 | 0 | 0 | 0 | 0 | 0 | 0 | 0 | 0 | 0 | 0 | 0 | 0 | 0 | 0 | 0 |
| | | detachment HS | 0 | 0 | 0 | 0 | 0 | 0 | 0 | 0 | 0 | 0 | 0 | 0 | 0 | 0 | 0 | 0 | 0 | 0 | 0 | 0 | 0 | 0 | 0 | 0 | 0 | 0 | 0 | 0 | 0 | 0 | 0 |
| | | Fracture NS | 0 | 0 | 0 | 0 | 0 | 0 | 0 | 0 | 0 | 0 | 0 | 0 | 0 | 0 | 0 | 0 | 0 | 0 | 0 | 0 | 0 | 0 | 0 | 0 | 0 | 0 | 0 | 0 | 0 | 0 | 0 |
| | | Fracture NA | 0 | 0 | 0 | 0 | 0 | 0 | 0 | 0 | 0 | 0 | 0 | 0 | 0 | 0 | 0 | 0 | 0 | 0 | 0 | 0 | 0 | 0 | 0 | 0 | 0 | 0 | 0 | 0 | 0 | 0 | 0 |
| | | Fracture centrum | 0 | 0 | 0 | 0 | 0 | 0 | 0 | 0 | 0 | 0 | 0 | 0 | 0 | 0 | 0 | 0 | 0 | 0 | 0 | 0 | 0 | 0 | 0 | 0 | 0 | 0 | 0 | 0 | 0 | 0 | 0 |
| | | Fracture HA | 0 | 0 | 0 | 0 | 0 | 0 | 0 | 0 | 0 | 0 | 0 | 0 | 0 | 0 | 0 | 0 | 0 | 0 | 0 | 0 | 0 | 0 | 0 | 0 | 0 | 0 | 0 | 0 | 0 | 0 | 0 |
| | | fracture HS | 0 | 0 | 0 | 0 | 0 | 0 | 0 | 0 | 0 | 0 | 0 | 0 | 0 | 0 | 0 | 0 | 0 | 0 | 0 | 0 | 0 | 0 | 0 | 0 | 0 | 0 | 0 | 0 | 0 | 0 | 0 |
| | | Extra intramem NS | 0 | 0 | 0 | 0 | 0 | 0 | 0 | 0 | 0 | 0 | 0 | 0 | 0 | 0 | 0 | 0 | 0 | 0 | 0 | 0 | 0 | 0 | 0 | 0 | 0 | 0 | 0 | 0 | 0 | 0 | 0 |
| | | Extra intramem NA | 0 | 0 | 0 | 0 | 0 | 0 | 0 | 0 | 0 | 0 | 0 | 0 | 0 | 0 | 0 | 0 | 0 | 0 | 0 | 0 | 0 | 0 | 0 | 0 | 0 | 0 | 0 | 0 | 0 | 0 | 0 |
| | | extra intramem centrum | 0 | 0 | 0 | 0 | 0 | 0 | 0 | 0 | 0 | 0 | 0 | 0 | 0 | 0 | 0 | 0 | 0 | 0 | 0 | 0 | 0 | 0 | 0 | 0 | 0 | 0 | 0 | 0 | 0 | 0 | 0 |
| | | Extra intramem HA | 0 | 0 | 0 | 0 | 0 | 0 | 0 | 0 | 0 | 0 | 0 | 0 | 0 | 0 | 0 | 0 | 0 | 0 | 0 | 0 | 0 | 0 | 0 | 0 | 0 | 0 | 0 | 0 | 0 | 0 | 0 |
| | | Extra intramem HS | 0 | 0 | 0 | 0 | 0 | 0 | 0 | 0 | 0 | 0 | 0 | 0 | 0 | 0 | 0 | 0 | 0 | 0 | 0 | 0 | 0 | 0 | 0 | 0 | 0 | 0 | 0 | 0 | 0 | 0 | 0 |
| | | lordosis | 0 | 0 | 0 | 0 | 0 | 0 | 0 | 0 | 0 | 0 | 0 | 0 | 0 | 0 | 0 | 0 | 0 | 0 | 0 | 0 | 0 | 0 | 0 | 0 | 0 | 0 | 0 | 0 | 0 | 0 | 0 |
| | | kyphosis | 0 | 0 | 0 | 0 | 0 | 0 | 0 | 0 | 0 | 0 | 0 | 0 | 0 | 0 | 0 | 0 | 0 | 0 | 0 | 0 | 0 | 0 | 0 | 0 | 0 | 0 | 0 | 0 | 0 | 0 | 0 |
| | | scoliosis | 0 | 0 | 0 | 0 | 0 | 0 | 0 | 0 | 0 | 0 | 0 | 0 | 0 | 0 | 0 | 0 | 0 | 0 | 0 | 0 | 0 | 0 | 0 | 0 | 0 | 0 | 0 | 0 | 0 | 0 | 0 |
| | | torsion | 0 | 0 | 0 | 0 | 0 | 0 | 0 | 0 | 0 | 0 | 0 | 0 | 0 | 0 | 0 | 0 | 0 | 0 | 0 | 0 | 0 | 0 | 0 | 0 | 0 | 0 | 0 | 0 | 0 | 0 | 0 |
| | | notochord mineral | 0 | 0 | 0 | 0 | 0 | 0 | 0 | 0 | 0 | 0 | 0 | 0 | 0 | 0 | 0 | 0 | 0 | 0 | 0 | 0 | 0 | 0 | 0 | 0 | 0 | 0 | 0 | 0 | 0 | 0 | 0 |
| | | IVL mineral | 0 | 0 | 0 | 0 | 0 | 0 | 0 | 0 | 0 | 0 | 0 | 0 | 0 | 0 | 0 | 0 | 0 | 0 | 0 | 0 | 0 | 0 | 0 | 0 | 0 | 0 | 0 | 0 | 0 | 0 | 0 |

**Table S2:** Parameters measured on the intervertabral space (IVS) transmission electron microscopy (TEM) images. Number of cells, and grading of key structures was taken directly from live images under the transmission electron microscope. Cells were counted in the IVL as shown on the schematic in Figure 6. The average number of different types of cells was calculated for both mutant and wild type siblings, respectively. For counting and measuring collagen layers, measuring bone thickness and measuring the angel of the IVL, images were taken and counts and measurements were taken from images.

| Parameter | Definition | Method |
|---|---|---|
| NUMBER OF CELLS | | |
| Number of chordoblasts | The number of chordoblasts present at the level of the intervertebral ligament were counted. | Cells were counted when the nucleus and/or the cell outline could be observed |
| Total number of cells in intervertebral space | Every cell that was present in the intervertebral region was counted. | |
| Cells on the mature collagen layer | The number of cells in contact with the outer surface of the mature collagen layer | |
| Cells on bone L and R | The sum of the number of cells on the left and right side on the bone was used. | |

| Thickness and number of collagen layers | | |
|---|---|---|
| Layers of mature collagen: | The mature collagen layer has plywood-like pattern in which the circumferentially and longitudinally oriented fibres can be recognised | The number of layers of circumferentially and longitudinally oriented fibres were counted on the TEM images |
| Mature collagen thickness: | The thickness of the mature collagen layer with the plywood-like pattern outside of the external elastic sheath of the notochord, | Measured using ImageJ |
| Mean thickness of the longitudinal layers | The thickness of the layers of longitudinally oriented collagen fibres in the mature collagen layer were measured and subsequently averaged | |
| Mean thickness of the circumferential layers: | The thickness of the layers of circumferentially oriented collagen fibres in the mature collagen layer were measured and subsequently averaged | |

| | | |
|---|---|---|
| Mean thickness of the vertebral endplate bone L and R: | The thickness of the bone of the vertebral endplate on the left and right side of the intervertebral ligament was measured and averaged between sides. The measure was taken perpendicular to the notochord sheath at the level of the vertebral endplate. | |
| grading of key structures | | |
| Angle | Generally, the intervertebral ligament shows symmetry when a central line is drawn through the middle of thickened elastin layer of the ligament to the center of the intervertebral disc. | The top of the middle line outside of the notochord was connected to the intersection of elastin layer and bone of the autocentrum touching the notochord sheath. Both angles left and right between the middle line and the connecting line were measured and their equalness was used as a measure of symmetry. |
| Elastin | A crescent shaped external elastic membrane is a typical feature in sections of the intervertebral disc. The elastin was graded from 0 to 3 depending on the severity of delamination and fragmentation. | 0: Crescent shaped external elastic membrane with/or no delamination with/or no fragmentation<br><br>1: Irregular shape of external elastic membrane with/or delamination with/or no fragmentation |

| | | |
|---|---|---|
| | | 2: Irregular shape of external elastic membrane with/or medium delamination with/or fragmentation<br><br>3: Unrecognisable shape of external elastic membrane with/or severe delamination with/or medium to severe fragmentation and/or absence of elastin |
| Autocentrum: | The autocentrum is the compact bone layer outside of the external elastic membrane of the notochord sheath at the level of the vertebral bone and is also continuous with the mature collagen layer of the intervertebral ligament. The plywood-like pattern present in the mature collagen layer is also recognisable in the autocentrum. The autocentrum was graded from 0 to 3 depending on how recognisable and disorganised the plywood-like pattern was. | 0: Autocentrum with plywood-like pattern recognisable and well organised as in the mature collagen layer of the intervertebral ligament<br><br>1: Autocentrum with plywood-like pattern recognisable but less well organised<br><br>2: Autocentrum recognisable but unorganised plywood-like collagen layer pattern<br><br>3: Autocentrum difficult to recognise and no organisation of the collagen fibres like a plywood-like pattern |

| | | |
|---|---|---|
| ER grade: | The endoplasmic reticulum (ER) of the cells in the intervertebral ligament, on the bone, mature collagen and in the immature collagen layers were graded from 0 to 3 depending on shape and size. | 0: Flat sheet-like cisternae with ribosomes on their border<br><br>1: Flat sheet-like cisternae interspersed with rounded ER cisternae<br><br>2: Many small or a few large rounded ER cisternae<br><br>3: Many large circular ER cisternae |

**Table S3:** Up and downregulated proteins in the three different OI mutant models, *col1a1a*$^{mh13/+}$, *col1a1a*$^{dc124/+}$ and *col1a2*$^{mh15/+}$, compared to their respective sibling wild-type controls. Proteins belonging to the zebrafish matrisome proteins (Nauroy et al., 2018) are highlighted in grey.

| ***Protein ID*** | ***Protein name*** | ***Gene name*** | ***Ensembl Gene ID*** |
|---|---|---|---|
| | | ***col1a1a***$^{mh13/+}$ | |
| A0A0R4ISS9 | protein C (inactivator of coagulation factors Va and VIIIa), a | proca | ENSDARG00000093079 |
| A5WWI5 | alpha-2-HS-glycoprotein 2 | ahsg2 | ENSDARG00000069293 |
| Q7ZVL5 | serpin peptidase inhibitor, clade E (nexin, plasminogen activator inhibitor type 1), member 2 | serpine2 | ENSDARG00000029353 |
| A0A0R4IR54 | Collagen, type XXII, alpha 1 | col22a1 | Not mapped |
| A0A2R8Q9B1 | Olfactomedin-like protein 3A; Olfactomedin-like domain-containing protein | olfml3a | ENSDARG00000061852 |
| A0A2R8QU81 | NEDD8 ubiquitin like modifier | nedd8;nedd8l | ENSDARG00000007989.7 |
| | | ***col1a1a***$^{dc124/+}$ | |
| R4GDT8 | si:ch211-132g1.3 | s+A10:F76i:ch211-132g1.3 | ENSDARG00000089477 |
| A2CEW3 | fibronectin 1b | fn1b | ENSDARG00000006526 |
| F1QV29 | complement component c3a, duplicate 2 | c3a.2 | ENSDARG00000087359 |
| B8A568 | myosin, heavy polypeptide 1.1, skeletal muscle | myhz1.1 | ENSDARG00000067990 |
| B8A565 | complement component 9 | c9 | ENSDARG00000016319 |
| F1R0Q1 | solute carrier family 2 member 1b | slc2a1b | ENSDARG00000007412 |
| B0S5W0 | collagen, type VIII, alpha 1a | col8a1a | ENSDARG00000077403 |
| A7YY10 | zgc:171772 | RPL37A | ENSDARG00000115271 |
| F1QQ04 | serpin peptidase inhibitor, clade H (heat shock protein 47), member 1b | serpinh1b | ENSDARG00000019949 |
| A0A0R4IXT8 | brevican | bcan | ENSDARG00000099412 |
| A0A0R4IXA9 | cyclase associated actin cytoskeleton regulatory protein 2 | cap2 | ENSDARG00000104478 |
| F1REF9 | proteoglycan 4b | prg4b | ENSDARG00000028163 |
| Q66I20 | serpin peptidase inhibitor, clade F (alpha-2 antiplasmin, pigment epithelium derived factor), member 1 | serpinf1 | ENSDARG00000069048 |
| | | ***col1a2***$^{mh15/+}$ | |
| Q568L5 | Aldo-keto reductase family 1 member A1-B | akr1a1b | ENSDARG00000052030.7 |
| A0A0R4IPV1 | aminolevulinate dehydratase | alad | ENSDARG00000100372 |
| Q7ZVB2; B0S7W5 | aldehyde dehydrogenase 9 family, member A1a, tandem duplicate 1; aldehyde dehydrogenase 9 family, member A1a, tandem duplicate 2 | aldh9a1a;aldh9a1a.2 | ENSDARG00000069100; ENSDARG00000055331 |

| | | | |
|---|---|---|---|
| F1R6R2 | Si:ch211-270n8.1 (Fragment) | apcs | ENSDARG00000045089.7 |
| Q7ZU89 | archain 1b | arcn1b | ENSDARG00000031214 |
| Q90485 | Hemoglobin subunit beta-2 | ba2l | ENSDART00000116849.4 |
| A0A0R4IXT8 | brevican | bcan | ENSDARG00000099412 |
| Q32LQ4 | Betaine--homocysteine S-methyltransferase 1 | bhmt | ENSDARG00000013430.10 |
| B3DHE9 | six-cysteine containing astacin protease 1 | c6ast1 | ENSDARG00000070314.6 |
| A0A2R8QRU4 | Caldesmon 1a | cald1a | ENSDARG00000070314.6 |
| B0S5W0 | collagen, type VIII, alpha 1a | col8a1a | ENSDARG00000077403 |
| Q1L8P0 | cartilage associated protein | crtap | ENSDARG00000018010 |
| F1R2S9 | extracellular matrix protein 2, female organ and adipocyte specific | ecm2 | ENSDARG00000071549 |
| A0A0R4IG09 | eukaryotic translation initiation factor 4h | eif4h | ENSDARG00000042252 |
| I3ISA6 | epiplakin 1 | eppk1 | ENSDARG00000096359 |
| Q1L8P1 | FKBP prolyl isomerase 9 | fkbp9 | ENSDARG00000005023 |
| A0A2R8QHG5 | Zgc:92027 | gca | ENSDARG00000020187.9 |
| Q5XIZ6 | glycerol-3-phosphate dehydrogenase 1 like | gpd1l | ENSDARG00000040024 |
| Q7T358 | G1 to S phase transition 1, like | gspt1l | ENSDARG00000098627 |
| F1QS28 | heterogeneous nuclear ribonucleoprotein A0a | hnrnpa0a | ENSDARG00000089302 |
| F1R2E0 | heterogeneous nuclear ribonucleoprotein M | hnrnpm | ENSDARG00000061735 |
| Q58EB2 | immunoglobulin like and fibronectin type III domain containing 1, tandem duplicate 1 | igfn1.1 | ENSDARG00000005526 |
| Q98862 | Indian hedgehog B protein;Hedgehog protein | ihhb;ihha | ENSDARG00000058815; ENSDARG00000058733 |
| Q6DHE6 | lysyl-tRNA synthetase 1 | kars1 | ENSDARG00000103799 |
| Q6NYL7 | keratin 18b | krt18b | ENSDARG00000028618 |
| F1R8U0 | keratin 94 | krt94;krt93;si:ch211-156l18.7 | ENSDARG00000044975 |
| Q6DHU3; F1QP28 | keratin 97; Keratin 96 (Fragment) | krt97; krt96 | ENSDARG00000000212 |
| Q90ZL2 | LanC antibiotic synthetase component C-like 1 (bacterial) | lancl1 | ENSDARG00000013741 |
| F1QSL3 | Galectin (Fragment) | lgals3b | ENSDARG00000044001.8 |
| F1QRV6 | nucleosome assembly protein 1-like 4b | nap1l4b | ENSDARG00000068868 |
| Q6NX06 | nucleobindin 2b | nucb2b | ENSDARG00000036291 |
| Q6NWY2 | oxidative stress responsive kinase 1b | oxsr1b | ENSDARG00000027500 |
| F1QSK1 | poly (ADP-ribose) polymerase family, member 3 | parp3 | ENSDARG00000003961 |
| Q7ZVK5 | poly(rC) binding protein 2 | pcbp2 | ENSDARG00000099039 |
| F1QP99 | protein disulfide isomerase family A, member 7 | pdia7 | ENSDARG00000014015 |

| | | | |
|---|---|---|---|
| A4U7F9 | procollagen-lysine, 2-oxoglutarate 5-dioxygenase 2 | plod2 | ENSDARG00000011821 |
| Q7ZTZ3 | phosphomannomutase 2 | pmm2;pmm1 | ENSDARG00000037654 |
| Q7ZUS7 | protein phosphatase 6, catalytic subunit | ppp6c | ENSDARG00000002949 |
| F1R3J9 | protein arginine methyltransferase 1 | prmt1;LOC556054;prmt8b | ENSDARG00000010246 |
| Q502Q7 | peripherin | prph | ENSDARG00000028306 |
| Q803I7 | phosphoserine aminotransferase 1 | psat1 | ENSDARG00000016733 |
| F1QGH9 | proteasome 26S subunit, non-ATPase 11b | psmd11b | ENSDARG00000005134 |
| A0A0R4IS70 | purine-rich element binding protein Ba | purba | ENSDARG00000068822 |
| Q7ZSZ0 | RAB1A, member RAS oncogene family b | rab1ab | ENSDARG00000029663 |
| F1Q7F8 | RNA binding motif protein 4.2 | rbm4.2 | ENSDARG00000055080 |
| Q6PBX9 | receptor accessory protein 5 | reep5 | ENSDARG00000100742 |
| Q7T399 | ras homolog family member Ca | rhoca | ENSDARG00000021309 |
| Q6P5K5 | ribosomal protein, large, P1 | rplp1 | ENSDARG00000021864 |
| Q6PBX2 | secretion associated, Ras related GTPase 1B | sar1b | ENSDARG00000103403 |
| Q7SXP0 | SEC22 homolog B, vesicle trafficking protein a and b | sec22bb;sec22ba | ENSDARG00000100435 |
| Q6PHD9 | Methanethiol oxidase | selenbp1 | ENSDARG00000024717.10 |
| Q7T309 | serpin peptidase inhibitor, clade B (ovalbumin), member 1, like 3 | serpinb1l3 | ENSDARG00000014556 |
| F1QQ04 | serpin peptidase inhibitor, clade H (heat shock protein 47), member 1b | serpinh1b | ENSDARG00000019949 |
| Q92008 | sonic hedgehog signaling molecule a | shha; shhb | ENSDARG00000068567 |
| Q6ZM17 | si:ch211-5k11.8 | si:ch211-5k11.8;hbae1.1;hbae5;hbae1.3;hbae3 | ENSDARG00000079078 |
| A3KNR2 | Novel protein | si:dkey-238c7.16 | ENSDARG00000111432.1 |
| F1Q7N8 | si:dkey-28b4.8 | si:dkey-28b4.8 | ENSDARG00000002840 |
| A0A2R8QEL5 | TAR DNA-binding protein 43 | tardbpa | ENSDARG00000004452.10 |
| A8DZJ0 | tubulin folding cofactor B | tbcb | ENSDARG00000068404 |
| Q6TNP9 | Tubulin alpha chain | tuba5 | ENSDARG00000042708.6 |
| F1R184 | thioredoxin domain containing 5 | txndc5 | ENSDARG00000009342 |
| F1Q5D8 | vitronectin b | vtna; vtnb | ENSDARG00000053831 |
| A0A0N4SU18 | Zgc:123103 (Fragment) | zgc:123103 | NSDARP00000135286 |

**Dataset S1 (separate file).** Completed template of the novel scoring matrix, designed to enable reproducible skeletal phenotyping based on Alizarin Red S mineral-stained whole skeletons.

**Dataset S2 (separate file).** List of all the identified proteins using liquid chromatography-tandem mass spectrometry (LC-MS/MS).

**Dataset S3 (separate file).** List of all the quantified proteins using liquid chromatography-tandem mass spectrometry (LC-MS/MS).

**Dataset S4 (separate file).** Differentially expression testing of quantified proteins by liquid chromatography-tandem mass spectrometry (LC-MS/MS). Protein expression levels were compared between each mutant and its respective WT sibling controls and significant differences were indicated with a '+' for upregulation and '-' for downregulation.
